# Soma-axon coupling configurations that enhance neuronal coincidence detection

**DOI:** 10.1101/405829

**Authors:** Joshua H Goldwyn, Michiel WH Remme, John Rinzel

**Affiliations:** Department of Mathematics and Statistics, Swarthmore College, Swarthmore, Pennsylvania, USA; Institute for Theoretical Biology, Humboldt University Berlin, Berlin, Germany; Center for Neural Science, New York University, New York, New York, USA; Courant Institute of Mathematical Sciences, New York University, New York, New York, USA

## Abstract

Coincidence detector neurons transmit timing information by responding preferentially to concurrent synaptic inputs. Principal cells of the medial superior olive (MSO) in the mammalian auditory brainstem are superb coincidence detectors. They encode sound source location with high temporal precision, distinguishing submillisecond timing differences among inputs. We investigate computationally how dynamic coupling between the “input” region (soma and dendrite) and the spike-generating “output” region (axon and axon initial segment) can enhance coincidence detection in MSO neurons. To do this, we formulate a two-compartment neuron model and characterize extensively coincidence detection sensitivity throughout a parameter space of coupling configurations. We focus on the interaction between coupling configuration and two currents that provide dynamic, voltage-gated, negative feedback in subthreshold voltage range: sodium current with rapid inactivation and low-threshold potassium current, *I*_*KLT*_. These currents reduce synaptic summation and can prevent spike generation unless inputs arrive with near simultaneity. We show that strong soma-to-axon coupling promotes the negative feedback effects of sodium inactivation and is, therefore, advantageous for coincidence detection. Furthermore, the “feedforward” combination of strong soma-to-axon coupling and weak axon-to-soma coupling enables spikes to be generated efficiently (few sodium channels needed) and with rapid recovery that enhances high-frequency coincidence detection. These observations detail the functional benefit of the strongly feedforward configuration that has been observed in physiological studies of MSO neurons. We find that *I*_*KLT*_ further enhances coincidence detection sensitivity, but with effects that depend on coupling configuration. For instance, in weakly-coupled models, *I*_*KLT*_ in the spike-generator compartment enhances coincidence detection more effectively than *I*_*KLT*_ in the input compartment. By using a minimal model of soma-to-axon coupling, we connect structure, dynamics, and computation. Here, we consider the particular case of MSO coincidence detectors. In principle, our method for creating and exploring a parameter space of two-compartment models can be applied to other neurons.

**Author summary:** Brain cells (neurons) are spatially extended structures. The locations at which neurons receive inputs and generate outputs are often distinct. We formulate and study a minimal mathematical model that describes the dynamical coupling between the input and output regions of a neuron. We construct our model to reflect known properties of neurons in the auditory brainstem that play an important role in our ability to locate sound sources. These neurons are known as “coincidence detectors” because they are most likely to respond when they receive simultaneous inputs. We use simulations to explore coincidence detection sensitivity throughout the parameter space of input-output coupling and to identify the coupling configurations that are best for neural coincidence detection. We find that strong forward coupling (from input region to output region), enhances coincidence detection sensitivity in our model and that low-threshold potassium current further improves coincidence detection. Our study is significant in that we detail how cell structure affects neuronal dynamics and, consequently, the ability of neurons to perform as temporally-precise coincidence detectors.

## Introduction

Neurons that spike selectively to multiple subthreshold inputs that arrive within brief time windows are coincidence detectors. Coincidence detection is a fundamental neural computation that allows the brain to extract information from the temporal patterns of synaptic inputs. In the cortex, neurons have biophysical specializations compatible with coincidence detection [1–7], but some have questioned whether temporally-precise computations are possible in cortex due to highly variable neural activity therein [8, 9]. In the early auditory pathway, the existence of coincidence detector neurons and their functional importance are widely valued [10–12]. Principal cells of the medial superior olive (MSO) in the mammalian auditory brainstem are a canonical example: they receive inputs originating from both ears [13, 14] and are sensitive to microsecond-scale differences in the timing of arriving inputs [15–18]. These coincidence detector neurons are critical for sound-source localization [19, for review] and likely play important roles in other aspects of binaural (“two-eared”) hearing such as sensitivity to interaural correlation [20, 21]

Temporally-precise neural coincidence detection requires specialized neural dynamics and circuitry. Coincidence detector neurons should have fast membrane dynamics with time-scales of integration shorter than the intervals between volleys of synaptic inputs [2, 22]. Inputs to coincidence detectors should also be brief and well-timed to precisely convey timing information. The requirements for effective coincidence detection in the auditory system are exceptionally stringent because auditory neurons must process inputs with temporal information at kilohertz-scale and higher [23–25]. Auditory brainstem circuitry is equipped with a suite of specializations to promote coincidence detection [26]. Afferent inputs to MSO cells are reliable and temporally-precise [27, 28], dendritic processing in MSO further enhances coincidence detection [25, 29–31], and voltage-gated currents that are partially active near resting voltage make MSO cells extremely fast and precise processors [25, 32]. Voltage-gated currents are also sources of dynamic negative feedback that contribute to the remarkable coincidence detection capabilities of these neurons. In MSO neurons, activation of low-threshold potassium (KLT) current and inactivation of sodium current are two identified sources of dynamic negative feedback [33–36]. In response to, say, a pair of brief excitatory inputs, these feedback mechanisms will transiently raise the spiking threshold after the first input, and thereby reduce the chance that the neuron will spike in response to the second input unless the inputs arrive nearly synchronously (within a coincidence detection “time window”).

We investigate the extent to which a “structural” specialization – namely, the coupling between the input region (soma and dendrite) and the output region (axon and axon initial segment) – can further optimize coincidence detection sensitivity. To do this, we develop a two-compartment neuron model as a minimal description of input-output coupling and systematically explore the effects of coupling configuration on coincidence detection sensitivity.

The two-compartment formulation is motivated by observations that spike generation in MSO likely occurs in the axon or axon initial segment [37, 38]. Furthermore, sodium channels in the soma are inactivated near resting potentials [39] and spikes are small and graded in the soma [37], suggesting the soma does not participate in spike generation. Indeed, an absence (or small amount) of sodium in the soma appears as a general design principle for temporally-precise auditory neurons [40]. Studies of coincidence detector cells in the avian auditory brainstem have shown that a passive soma can enhance coincidence detection [41] and that the distribution of voltage-gated channels in the axon initial segment undergoes activity-dependent modulation [42, 43] to improve coincidence detection, perhaps in an optimal manner [44]. A two-compartment formulation neglects the helpful contributions of dendritic processing to coincidence detection, but the role of dendrites has been considered in detail in previous studies [25, 29–31].

In this study, we systematically relate forward coupling (soma-to-axon) and backward coupling (axon-to-soma) strengths to model parameters. We explore this parameter space and find the coupling configurations that enhance coincidence detection sensitivity. Specifically, we identify strong soma-to-axon coupling as a “natural” configuration for neural coincidence detection because it engages sodium inactivation as a mechanism that transiently increases spike threshold on the time-scale of synaptic inputs and prevents firing to inputs that do not arrive concurrently. Moreover, the combination of strong soma-to-axon and weak axon-to-soma coupling generate spikes more efficiently (requires fewer sodium channels) and with shorter refractory periods than other models. This “feedforward” configuration enhances high-frequency coincidence detection and represents distinct advantages over one-compartment “point neuron” models that cannot exhibit this asymmetric coupling configuration. We observe that KLT current provides additional benefits for coincidence detection sensitivity, but these benefits depend on coupling configuration and where KLT current is located in the two-compartment structure. For instance, coincidence detection sensitivity in neurons with weak soma-to-axon coupling can be substantially improved if KLT current is co-localized with spike-generating currents.

We select passive properties to match known physiological characteristics of MSO neurons, so our observations apply directly to those canonical coincidence detectors. Nonetheless, our method for systematically exploring the parameter space of coupling configurations can be applied to study the relationships between structure, dynamics, and computation in other neurons that are well-described by a two-compartment idealization [24, 45–50]. In particular, a useful aspect of our work is that we show how to explicitly construct two-compartment models that satisfy the constraint of having (nearly) identical passive dynamics in the input compartment.

## Materials and methods

### Two-compartment model parameterized by coupling strength

We construct and analyze a minimal description of a neuron that separates the input region (soma and dendrites) from the spike generating region (axon and axon initial segment) of a cell. This “two-compartment model” [45] has the form:

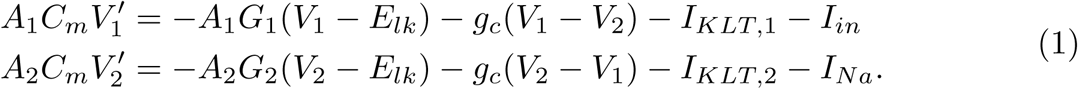

The dynamical variables *V*_*i*_ (*i* = 1, 2) describe the membrane potential in each compartment. Passive parameters in the model are membrane capacitance per area (*C*_*m*_), axial conductance (*g*_*c*_), reversal potential of leak current (*E*_*lk*_), compartment surface area (*A*_*i*_), and membrane leak conductance density (*G*_*i*_). Parameters subscripted with *i* can take different values in the two compartments (*i* = 1, 2). To simplify notation, we will often omit the explicit reference to membrane area and instead use the notation *c*_*i*_ = *A*_*i*_*C*_*m*_ and *g*_*i*_ = *A*_*i*_*G*_*i*_ for *i* = 1, 2. The first compartment (abbreviation: Cpt1) receives input current (*I*_*in*_) and the second compartment (abbreviation: Cpt2) is the site of spike-generating sodium current (*I*_*Na*_). In some simulations we also include dynamic (voltage-gated) low-threshold potassium current (*I*_*KLT*_). These currents are described in more detail below.

We use standard neurophysiological measures of passive activity in the soma to determine some parameters, and vary other parameters to create a “family” of two-compartment models distributed in a two-dimensional parameter space. We select model parameters so that, regardless of coupling configuration, the passive dynamics in Cpt1 are nearly identical regardless of the strength of coupling between compartments. This novel formulation allows us to meaningfully and systematically probe the dynamics of the model. The two parameters that define coupling configuration are introduced below. They describe strength of “forward” coupling (Cpt1 to Cpt2) and “backward” coupling (Cpt2 to Cpt1).

The properties we match to experimental measurements include resting potential in the soma (*V*_*rest*_), input resistance for input to the soma (*R*_*in*_), and exponential time constant (*τ*_*exp*_) with which soma voltage returns to rest following a brief perturbation. We use the following values based on in vitro measurements of gerbil MSO neurons [32]: *V*_*rest*_ = -58 mV, *R*_*in*_ = 8.5 MΩ, and *τ*_*exp*_ = 340 *µ*s. We first match these properties using a model with passive dynamics (by setting *I*_*Na*_ and *I*_*KLT*_ to zero). After identifying parameter relations that satisfy these constraints, we discuss how to introduce sodium and KLT currents.

In the passive model, the resting potential is identical to the reversal potential and we have *V*_*rest*_ = *E*_*lk*_. We now determine the remaining parameters based on the values of *R*_*in*_ and *τ*_*exp*_. It is convenient to rescale the voltage equations by *g*_*i*_ + *g*_*c*_ and to introduce terms that represent the deviation of voltage from rest: *U*_*i*_ = *V*_*i*_ –*E*_*lk*_ (for *i* = 1, 2). This yields the following equations for passive and subthreshold dynamics (*I*_*Na*_ and *I*_*KLT*_ removed, for now):

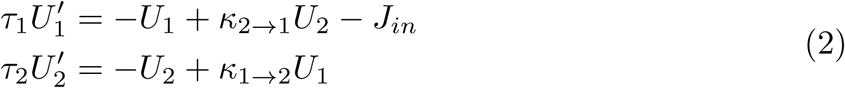

The rescaled input current is denoted *J*_*in*_ = *I*_*in*_*/*(*g*_1_ + *g*_*c*_). The time constants *τ*_*i*_ = *c*_*i*_*/*(*g*_*i*_ + *g*_*c*_) describe the passive dynamics of the *i*^*th*^ compartment (*i* = 1, 2) when the other compartment is held at its resting voltage.

We have also introduced in Eq. 2 the two parameters that describe coupling strength. The forward coupling parameter is *κ*_1*→*2_. Formally, it is the ratio of voltages *U*_2_*/U*_1_ at steady state in response to a constant current applied to Cpt1. Similarly, the backward coupling parameter is *κ*_2*→*1_. It is the ratio *U*_1_*/U*_2_ at steady state in response to constant current applied to Cpt2. These quantities are attenuation factors that take values between zero (complete attenuation) and one (no attenuation). We find it more intuitive to refer to these constants as measures of coupling strength – values near zero represent weak coupling, values near one indicate strong coupling. We will refer throughout to *κ*_1*→*2_ as the forward coupling parameter (“soma-to-axon” coupling) and *κ*_2*→*1_ as the backward coupling parameter (“axon-to-soma” coupling). The relationship between coupling parameters and conductance parameters in Eq. 1 are:

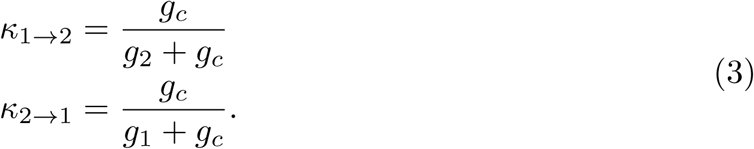

Next, we will show how to invert these equations to simply and uniquely define all passive model parameters for any combination of *κ*_1*→*2_ and *κ*_2*→*1_. We only require prior knowledge of *R*_*in*_ and *τ*_*exp*_ (experimentally measurable parameters) and the assumption that the area of Cpt1 is much larger than the area of Cpt2. We denote the ratio of compartment areas as *α* = *A*_2_*/A*_1_. We will always use *α* = 0.01 in this study. This assumption is plausible for cells with input regions that are much larger than spike-generating regions, and is consistent with previous models of auditory coincidence detector neurons [24, 38].

We find the axial conductance (*g*_*c*_) by expressing it in terms of input resistance and the coupling coefficients. By setting 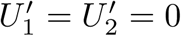 in Eq. 2, we find the steady state relations 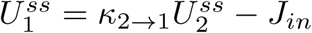 and 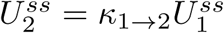. From these relations, and the steady-state input resistance (applying Ohm’s Law), it follows that 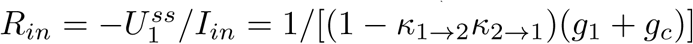. Solving for *g*_*c*_ and after some substitutions we find

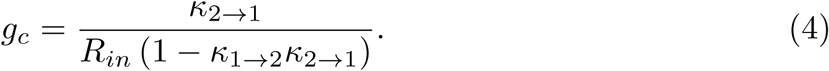

The remaining parameters determine the passive dynamics of *V*_1_, so they depend on *τ*_*exp*_. In a one-compartment passive model, *τ*_*exp*_ is identical to the membrane time constant and its value is *c*_1_*/g*_1_. In a two-compartment model, coupling between the compartments introduces a second time-scale that can influence the rate at which *V*_1_ returns to rest after a brief perturbation (Rall’s equalization time constant [51], and see also [52]). Some additional analysis is required, therefore, to relate *τ*_*exp*_ to model parameters.

We invoke the assumption that the input region is much larger than the output region (*A*_1_ *» A*_2_, or, equivalently, take 0 *< α «* 1) and observe that this can create a separation of time-scales in the passive dynamics of *U*_1_ and *U*_2_. The ratio of time constants in the two compartments is *τ*_2_*/τ*_1_ = (*c*_2_*/c*_1_)[(*g*_1_ + *g*_*c*_)*/*(*g*_2_ + *g*_*c*_)]. After some substitutions, and using the assumption that *C*_*m*_ is identical in both compartments, we find that

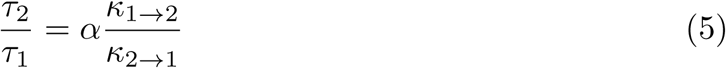

We restrict ourselves to coupling configurations for which *κ*_1*→*2_*/κ*_2*→*1_ does not exceed ten, so that *τ*_1_ is an order of magnitude larger than *τ*_2_ (recall we use α = 0.01). In this scenario, we can segregate the passive dynamics into a slow variable (*U*_1_) and a fast variable (*U*_2_). The ratio of time-constants is a small parameter which we denote by 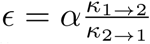. For *ε* close to zero, we can make the approximation that *U*_2_ evolves “instantaneously” (on the fast time-scale) to its *U*_1_-dependent steady-state value of 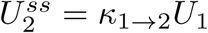. On the slow-time scale, *U*_2_ takes this instantaneous value and the dynamics of *U*_1_ are (to leading order in the small parameter *ε*):

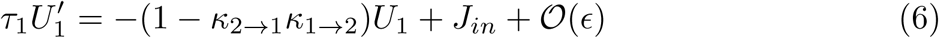

In other words, in cases when *τ*_1_ ≫ *τ*_2_, the passive dynamics in Cpt1 are approximately linear and one can match the time-scale of *U*_1_ to the experimentally-observed membrane decay time by setting *τ*_1_ = *τ*_*exp*_(1 − *κ*_1*→*2_*κ*_2*→*1_). This “slaving” of *U*_2_ to *U*_1_ is valid for describing passive subthreshold dynamics. If sodium current is included, then the dynamics are non-linear and spike-generation is possible. On the slow time-scale, spike-generation is represented by a discontinuous “jump” to a fixed point at higher values of *U*_1_ and *U*_2_.

To summarize our method: we use values for three standard neurophysiological measures of (passive) soma dynamics (*R*_*in*_, *τ*_*exp*_ and *E*_*lk*_), we choose *α* (the ratio of surface areas *A*_2_*/A*_1_) to be small (*α* = 0.01 in all simulations), and we let the two coupling constants define a two-dimensional parameter space of soma-axon coupling. For any coupling configuration, we can then uniquely determine the passive parameters in Eq. 1. The parameter relationships, described above, are:

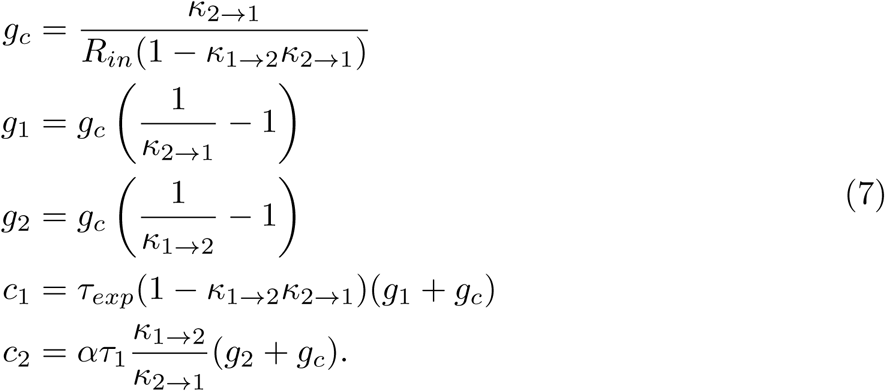

Using these parameter relations guarantees that passive dynamics in Cpt1 remain nearly identical as we explore neural dynamics and coincidence detection sensitivity in the two-dimensional parameter space of coupling strengths.

We only consider coupling configurations in which forward coupling is stronger than backward coupling (*κ*_1*→*2_ *≥* *κ*_2*→*1_). This corresponds to an assumption that signals propagate forward from the soma to the axon with less attenuation than signals that backpropagate from the axon to the soma. This condition is appropriate for MSO neurons, since in vitro recordings show weak backpropagation of action potentials to the soma and dendrites [31, 37].

### Low-threshold potassium model

A voltage-gated low-threshold potassium current (*I*_*KLT*_) is prominent in MSO neurons and thought to improve coincidence detection sensitivity [53]. We model *I*_*KLT*_ in the *i*^*th*^ compartment with the equation

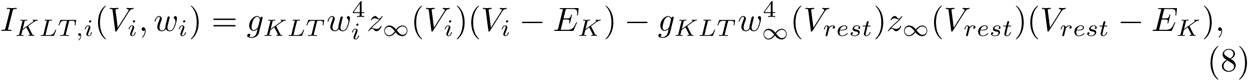

where the reversal potential is *E*_*K*_ = −106 mV. We include the second term so that the addition of KLT current does not alter the resting potential (*I*_*KLT,i*_ = 0 when *V*_*i*_ = *V*_*rest*_, for *i* = 1 or 2.). Equivalently, one could adjust *E*_*lk*_ to counterbalance the amount of KLT current active at rest, and omit this correction term. The dynamics of the activation variable *w*_*i*_ are as in [32] at a temperature of 35°C:

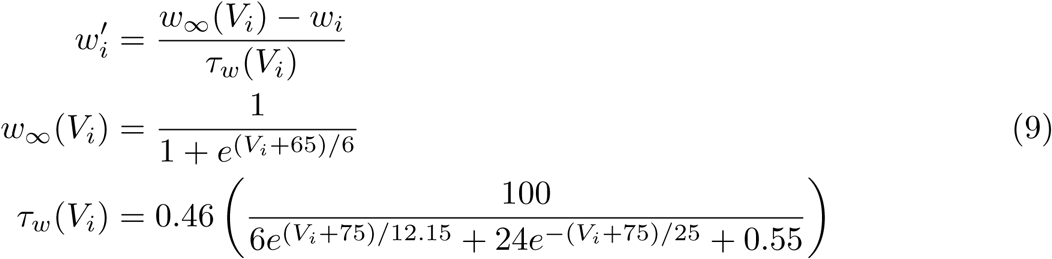

The inactivation variable is slow (time-scale of several hundred milliseconds), so we make the simplification that its value is fixed at the steady state *z*_*∞*_(*V*_*rest*_) where the steady state function is [32]

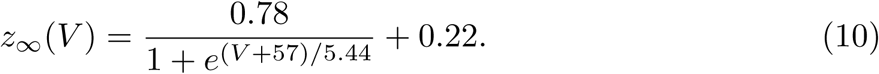

Next we discuss how we include dynamic *I*_*KLT*_ in the two-compartment model. We omit subscripts, for ease of presentation, but the same method applies for dynamic KLT current in either compartment. We first find the passive leak conductance in the relevant compartment using Eq. 7. Call this *g*_*lk*_. We then reduce this conductance by some amount, typically 10%. In other words, we set the leak conductance in the relevant compartment (*g*_*i*_) to 0.9*g*_*lk*_. Lastly, we set *g*_*KLT*_ to the value that preserves the total conductance in the compartment at the resting potential. In some simulations, we leave the KLT conductance fixed at its resting value. We refer to this case as “frozen” KLT – the KLT current acts as a leak current and the subthreshold dynamics are the same as the original passive model. In other simulations, we allow KLT conductance to depend on voltage. We refer to this case as “dynamic” KLT. To include dynamic KLT in Cpt1, for example, we would choose *g*_*KLT*_ so that it satisfies the equation 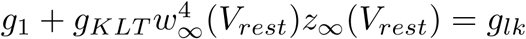 and allow the KLT activation variable *w* to evolve according to Eq. 9.

### Sodium current model

The second compartment represents regions of the cell in which spikes are generated, presumably the axon initial segment or other excitable regions in the axon [38]. We use a reduced model of sodium current, adapted from earlier models of auditory brainstem neurons [32, 54], to produce spikes:

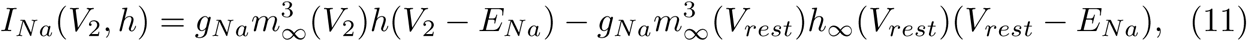

where the sodium reversal potential is *E*_*Na*_ = 55 mV. The second term is included so that *I*_*Na*_ = 0 when *V*_2_ is at its resting value. Equivalently, one could adjust *E*_*lk*_ to counterbalance the amount of sodium current active at rest, and omit this correction term. Setting *I*_*Na*_ = 0 at rest simplifies analysis of the model and is appropriate for MSO neurons since most sodium channels are inactivated for membrane potential near rest [39]. We assume that activation of sodium is sufficiently fast to justify the approximation that the gating variable *m* instantaneously reaches its voltage-dependent equilibrium value *m*_*∞*_(*V*_2_) [34]. The gating variable *h* governs inactivation of the sodium current and has dynamics as in [32]:

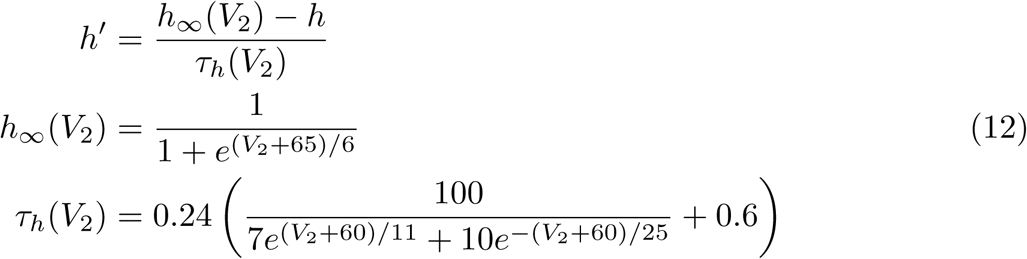

These are the same as in [54], but with temperature adjusted to 35°C, and also note the resting membrane potential in our model is -58 mV as opposed to -65 mV in [54]

Our primary objective is to determine the effects of coupling configuration on coincidence detection. Maximal conductance *g*_*Na*_ determines excitability and spike threshold in the model neuron and thus also influences coincidence detection sensitivity [22]. Rather than setting *g*_*Na*_ to an arbitrarily chosen value, we explore a range of *g*_*Na*_ to determine the best possible coincidence detection sensitivity over this range of *g*_*Na*_, for each coupling configuration. We explain our method for choosing *g*_*Na*_ values in more detail below.

### Synaptic input model

We generate synaptic inputs to the two-compartment model using a model of the auditory periphery [55]. This model includes the effects of cochlear filtering and nonlinearities, inner hair cell activity, and synaptic transmission, and generates auditory nerve spike trains. As inputs to this model, we use sine waves that represent pure tone sounds. We perform simulations with frequencies ranging from 200 Hz to 700 Hz at a level of 70 dB. The neuron model receives two streams of auditory nerve inputs representing (conceptually) inputs from the two ears, see the schematic in Fig. 1. The sine waves to the two “ears” are presented either with identical timing to generate “coincident” inputs, or with a time delay to simulate “non-coincident” inputs.

**Fig 1.**
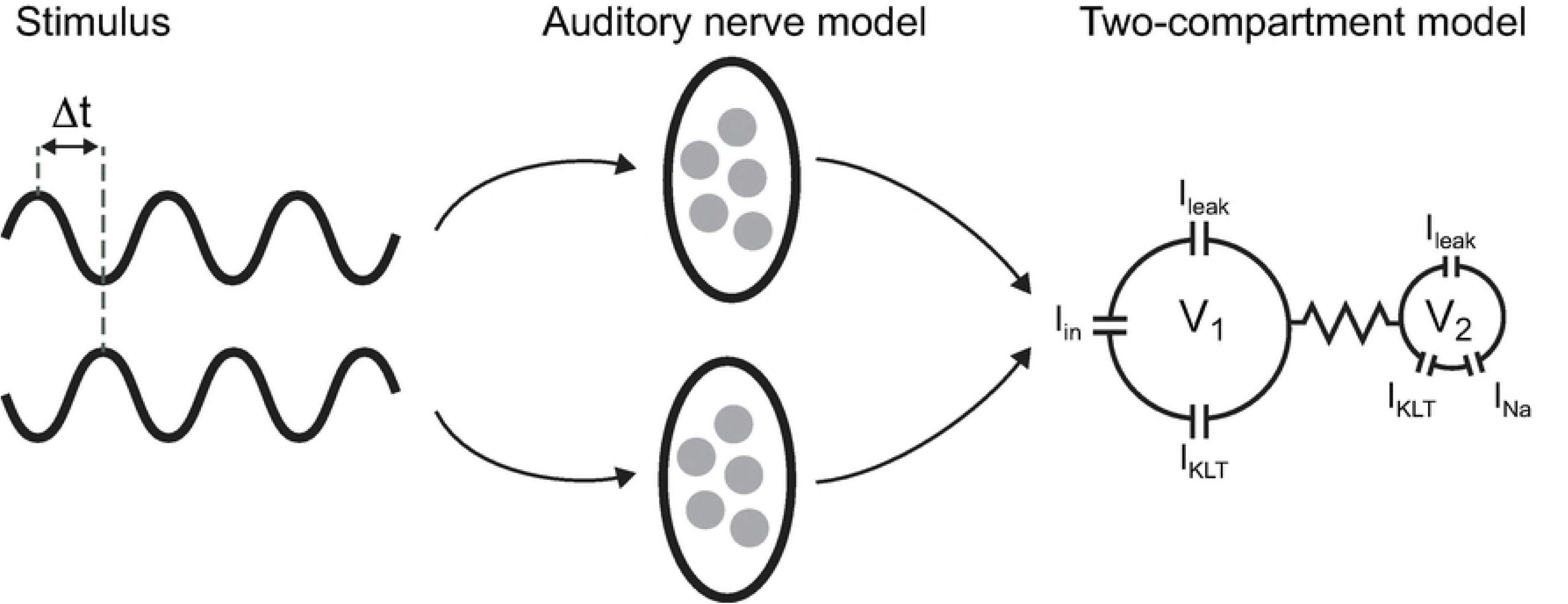
Schematic of simulations to test coincidence detection sensitivity of two-compartment neuron model. Stimulus is a pair of sine-wave inputs (frequencies range from 200 Hz to 700 Hz) that are presented either in phase with one another (“coincident inputs”) or with a time delay (“non-coincident inputs”, indicated by Δ*t* in this figure). Sine-wave inputs are delivered to a model of the auditory periphery [55]. We extract from this model simulated spiking responses of five auditory nerve fibers for each sine-wave. These simulated spike trains are used to create excitatory current (*I*_*in*_ in this figure) that is delivered to Cpt1.

MSO coincidence detectors receive a small number of synaptic inputs [56], so we use the auditory nerve model to simulate five independent input sequences of spike times per “ear.” Each auditory nerve spike time creates an excitatory post-synaptic conductance (EPSG) described by a double exponential function [18]:

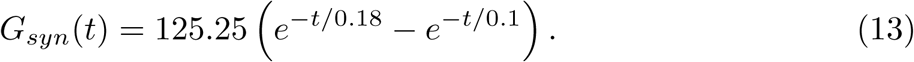

and these EPSGs are transformed into synaptic current (EPSCs) according to the equation

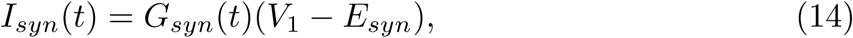

where the reversal potential for the excitatory current is *E*_*syn*_ = 0 mV. We set the constant scaling factor in the definition of *G*_*syn*_(*t*) (Eq. 13) so that a single excitatory input depolarizes *V*_1_ by roughly 6 mV, a value consistent with measurements of MSO neurons’ responses to synaptic excitation in vitro [57].

We view this as an adequate input for probing coincidence detection in a MSO-like two-compartment model using quasi-realistic stimuli. It is not meant to be a “complete” description of neural processing in the MSO-pathway. Notably, we do not include spherical bushy cells in the cochlear nucleus that may enhance temporal precision of afferent inputs to MSO neurons [27], nor do we include inhibitory inputs that appear to modify time-difference tuning of MSO coincidence detectors [57–63], but see also [18].

In some simulations, as we will make clear in the context of the Results, we use more simplistic inputs such as steps or ramps of current injected directly into the input compartment to study response characteristics of the model.

### Measure of coincidence detection sensitivity

We measure firing rate (spikes per second) generated by the two-compartment model in response to “coincident” inputs (identical sine wave stimuli to the two “ears” of the auditory nerve model) and “non-coincident” inputs. In some simulations we generate “non-coincident” inputs by using sine wave stimuli that were anti-phase to the two “ears” of the auditory nerve model. For example, for a 500 Hz stimuli, the two sine waves would have a 1 ms time difference. In this construction of non-coincident inputs, the time difference between the sine waves shortens with increasing frequency. To confirm that this dependence of time difference on frequency does not bias our results, we also perform simulations in which we generate non-coincident inputs by using sine wave stimuli to the auditory nerve model with a fixed time difference of 500 *µ*s.

To measure coincidence detection sensitivity, we compute the difference in firing rates for responses to coincident and non-coincident inputs. We compute firing rates (spikes per second) by counting the number of spikes generated in trials of length 250 ms. We then calculate mean and standard error of firing rates from 100 repeated trials. Large-scale calculations to sweep over parameter space were performed using Matlab simulation code executed on computers managed by the Ohio Supercomputer Center. The ordinary differential equations defining the two-compartment model were solved numerically using the Matlab command *ode15s* (a variable-step, variable-method solver useful for stiff systems). Simulation code is available at https://github.com/jhgoldwyn/TwoCompartmentModel.

A “good” coincidence detector neuron would be one with a large difference in firing rates for these two conditions. Firing rate difference measures of coincidence detection sensitivity have been used in related studies [64, 65]. Other measures have been considered, including Fisher information [38], width of time-difference tuning curves [62], and quality factor (similar to d-prime) [22]. The “right” measurement of coincidence detection sensitivity remains, as these alternatives reveal, an open question (and one wrapped up in ongoing debates regarding the nature of the neural code for sound source location [67]).

One justification for comparing in-phase to out-of-phase firing rates is that it is relevant to a system that uses a “two channel” representation of auditory space in which sound location is represented by the difference in firing rates between two populations of cells tuned to distinct time-differences [66]. We suggest an additional perspective based on an analogy to signal classification theory and the receiver operating characteristic (ROC) [68].

Consider a coincidence detector neuron responding to a periodic (sine wave) stimulus. Each cycle of the stimulus evokes a volley of synaptic inputs that may or may not be temporally aligned with one another. The task of the coincidence detector neuron is to respond (generate a spike) if the synaptic inputs arrive within a brief time window and to not respond (not spike) if the synaptic inputs are dispersed in time. From this perspective, a coincidence detector neuron is an “observer” of its own synaptic inputs and it signals the presence of coincident inputs by generating a spike. Chance [69] has articulated a similar approach for measuring synaptic efficacy.

Extending the analogy, for each two-compartment model (parameterized by coupling configuration), we construct ROC curves by plotting “hit” rate (firing rate to coincident inputs) against “false alarm” rate (firing rate to non-coincident inputs) for varying values of the sodium conductance *g*_*Na*_. Sodium conductance controls the overall excitability of the model and operates as the threshold parameter in ROC analysis. To compare coincidence detection sensitivity across coupling configurations, we simulate the model for a range of *g*_*Na*_ values and define coincidence detection sensitivity to be the maximum firing rate difference. In this way, we identify the *g*_*Na*_ level for which the neuron, acting as an observer of its inputs, is the best possible coincidence detector. In other words, we identify the *g*_*Na*_ value that maximizes “hits” (spikes generated in response to coincident inputs) while minimizing “false alarms” (spikes generated in response non-coincident inputs), for a given coupling configuration and stimulus. Fig. 2A illustrates this calculation (illustration only, not actual simulation data).

**Fig 2.**
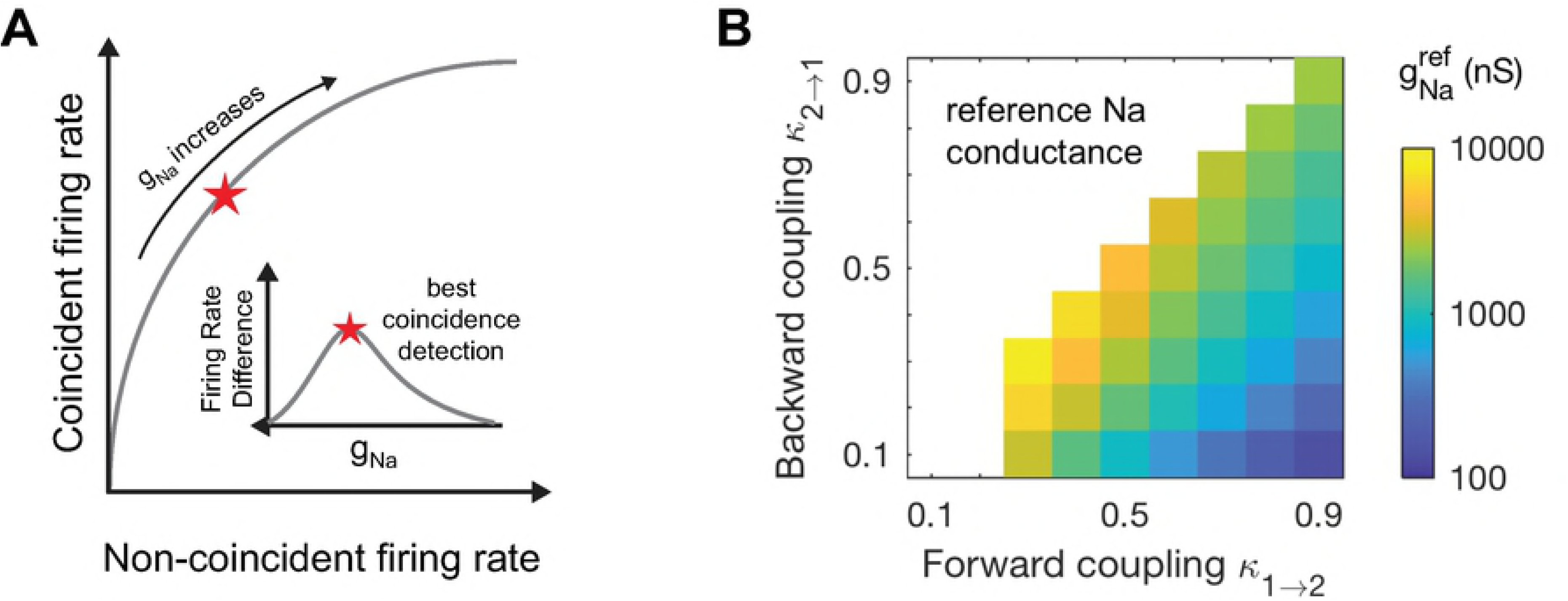
Coincidence detection sensitivity measurement and 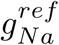 values. A. Firing rate to coincident inputs and non-coincident inputs increase with *g*_*Na*_ (cartoon, not actual data). We sweep across a range of *g*_*Na*_ values and quantify coincidence detection sensitivity as the maximum firing rate difference across all *g*_*Na*_ values used (inset). We draw an analogy to the signal receiver operating characteristic (ROC) curve: coincident firing rate is the “hit” rate, non-coincident firing rate is the “false alarm” rate, and *g*_*Na*_ sets the detection threshold. This panel is for illustration only and does not portray actual simulation data. **B:** Reference values for sodium conductance (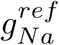; the smallest value of *g*_*Na*_ at which a pair of simultaneous EPSG events evokes a spike).

The *g*_*Na*_ values that produced similar spiking activity for different coupling configurations could vary by orders of magnitude, so we required a reasonable way to determine the appropriate range of *g*_*Na*_ values to use. For each coupling configuration, we set a reference value for *g*_*Na*_, denoted 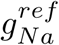, by finding the smallest *g*_*Na*_ value for which a pair of coincident EPSGs could evoke a spike. The rationale for this definition of 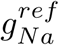 is that, for any *g*_*Na*_ value larger than 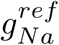, the model neuron could possibly spike in response to two non-coincident EPSGs. For *g*_*Na*_ significantly larger than 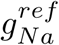, the model neuron could even spike in response to a single EPSG. By restricting *g*_*Na*_ to values near 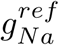, we ensured that the model neurons were operating as coincidence detectors and not responding to multiple inputs that lacked temporal alignment. Values of 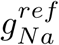 in the parameter space of forward and backward coupling strengths are shown in Fig. 2B. Values of 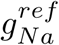 increase if dynamic KLT current is present in the model. Replacing 10% of *g*_*lk*_ with KLT conductance increased reference *g*_*Na*_ values by 5% to 30% higher depending on coupling configuration, see S1 Fig.

We then measure coincidence detection sensitivity using *g*_*Na*_ values that ranged from 0.2 to 2.2 times 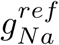, in increments of 0.05 times 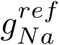. From the firing rate differences measured across this range of *g*_*Na*_ values, we identify the maximum firing rate difference and use this as our measure of coincidence detection sensitivity. As a result of this process, we obtain different “best” *g*_*Na*_ values depending on input frequency, coupling configuration, and KLT currents. An implication of this approach is that neurons should modulate sodium conductance based on stimulus parameters and physiological conditions. We do not pursue this idea here, but see [42, 44] for relevant studies and S2 Fig for supporting results showing changes of “best” *g*_*Na*_ with stimulus frequency.

## Results

### Parameterization of a family of two-compartment models by coupling strength

As discussed in Materials and methods, the forward coupling parameter (soma-to-axon, *κ*_1*→*2_) and backward coupling parameter (axon-to-soma, *κ*_2*→*1_) characterize the passive dynamics of a two-compartment model (see Eq. 3). By defining model parameters according to Eq. 7, we construct a family of models with (nearly) identical passive dynamics in Cpt1. Recall we select model parameters based on reported properties of MSO neurons: steady state input resistance is 8.5 MΩ, the decay time constant is 340 *µ*s, and the resting potential is –58 mV [32]. We also assume the capacitance per unit area is the same in both compartments and that the surface area of Cpt1 was 100 times larger than the surface area of Cpt2 (*α* = 0.01). Using a typical value of membrane capacitance *C*_*m*_ = 0.9 pF/cm^2^ [70], we find that capacitance in Cpt1 is *C*_1_ = 40 pF. The area of Cpt1 is *A*_1_ = 4444 *µ*m^2^, which is a plausible value for the surface area of the soma and dendrite regions of an MSO neuron [31, 71].

In order to maintain identical passive dynamics in Cpt1 across coupling configurations, the leak conductance (*g*_1_, *g*_2_) and axial conductance (*g*_*c*_) vary with the values of the coupling parameters as shown in Fig. 3. Leak conductance in Cpt1 increases as forward coupling strength increases but decreases as backward coupling strength increases (Fig. 3A). Leak conductance in Cpt2 exhibits the opposite dependence on coupling strength: *g*_2_ decreases as forward coupling strength increases and increases as backward coupling strength increases (Fig. 3B). The axial conductance connecting the two compartments depends primarily on the strength of backward coupling – the contour lines in Fig. 3C are nearly horizontal except in cases of strong forward and backward coupling (upper right corner). The upper left in each panel of Fig. 3 is empty because we only consider coupling configurations for which forward coupling is not weaker than backward coupling (*κ*_1*→*2_ *≥* *κ*_2*→*1_).

**Fig 3.**
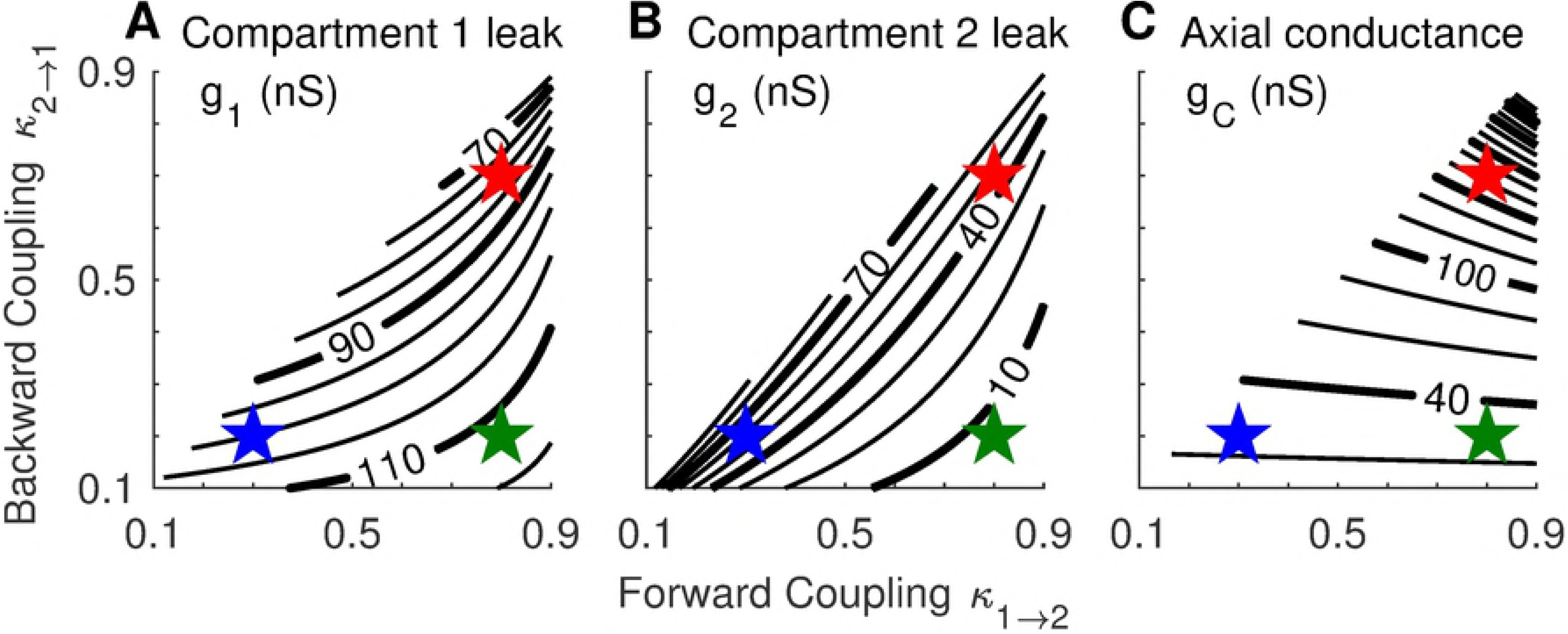
Passive parameters as a function of forward and backward coupling strength. Parameter values derived using Eq. 7 and the assumptions that the decay time constant for *V*_1_ is *τ*_*exp*_ = 0.34 ms, input resistance in Cpt1 is *R*_*in*_ = 8.5 MΩ, and the surface area of Cpt1 is one hundred times larger than the surface area of Cpt2. The upper left half of the parameter space is empty because we did not consider models for which forward coupling was weaker than backward coupling. Colored stars mark the locations of three configurations we examine in detail below: the weakly-coupled model (*κ*_1*→*2_ = 0.3, *κ*_2*→*1_ = 0.2; blue), the forward-coupled model (*κ*_1*→*2_ = 0.8, *κ*_2*→*1_ = 0.2; green), and the strongly-coupled model (*κ*_1*→*2_ = 0.8, *κ*_2*→*1_ = 0.7; red).

To explore how coupling configuration modifies neural dynamics, we will often compare three models near the edges of the coupling parameter space. These are a *weakly-coupled model* (*κ*_1*→*2_ = 0.3, *κ*_2*→*1_ = 0.2), a *strongly-coupled model* (*κ*_1*→*2_ = 0.8, *κ*_2*→*1_ = 0.7), and a *forward-coupled model* (*κ*_1*→*2_ = 0.8, *κ*_2*→*1_ = 0.2). The locations of these three models in the coupling parameter space are shown as colored stars in Fig. 3. We remark that “complete” coupling (*κ*_1*→*2_ = *κ*_2*→*1_ = 1) is equivalent to a one-compartment “point” neuron model because voltages in the two compartments are the same in this case.

### Passive dynamics in first compartment are nearly identical across coupling configurations

Our parameterization method is designed to maintain the same voltage response in Cpt1 (*V*_1_) regardless of the coupling configuration. In fact, due to the strong separation of time scales between the two compartments (recall Eq. 5), the voltage in Cpt1 is governed approximately by linear dynamics with time constant *τ*_*exp*_ (see Eq. 6) and the voltage in Cpt2 is *V*_2_ ≈ *E*_*lk*_ + *κ*_1*→*2_(*V*_1_ –*E*_*lk*_). These approximations are valid to leading order in the small parameter 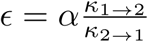. We remind the reader that these calculations are performed in the case of passive dynamics — i.e. for a model without spike-generating sodium current (*g*_*Na*_ = 0) and with “frozen” low-threshold potassium current (*I*_*KLT*_ acts as a leak current, and in fact is equivalent to *g*_*KLT*_ = 0, see Materials and methods).

Simulations of passive two-compartment models illustrate how the parameterization method results in models with nearly identical *V*_1_ dynamics (Fig. 4B). We use darker colors to show time-courses of voltage in Cpt1 (*V*_1_) and lighter colors to show time-courses of voltage in Cpt2 (*V*_2_). *V*_1_ responses are nearly identical regardless of coupling configuration and attenuation of *V*_2_ responses depends on *κ*_1*→*2_. Time-courses of *V*_1_ and *V*_2_ shown here are responses to 500 Hz coincident inputs. The three coupling configurations in this figure are indicated in the schematic in the top row (from left to right in Fig. 4A: weakly-coupled, forward-coupled, and strongly-coupled, as defined previously).

**Fig 4.**
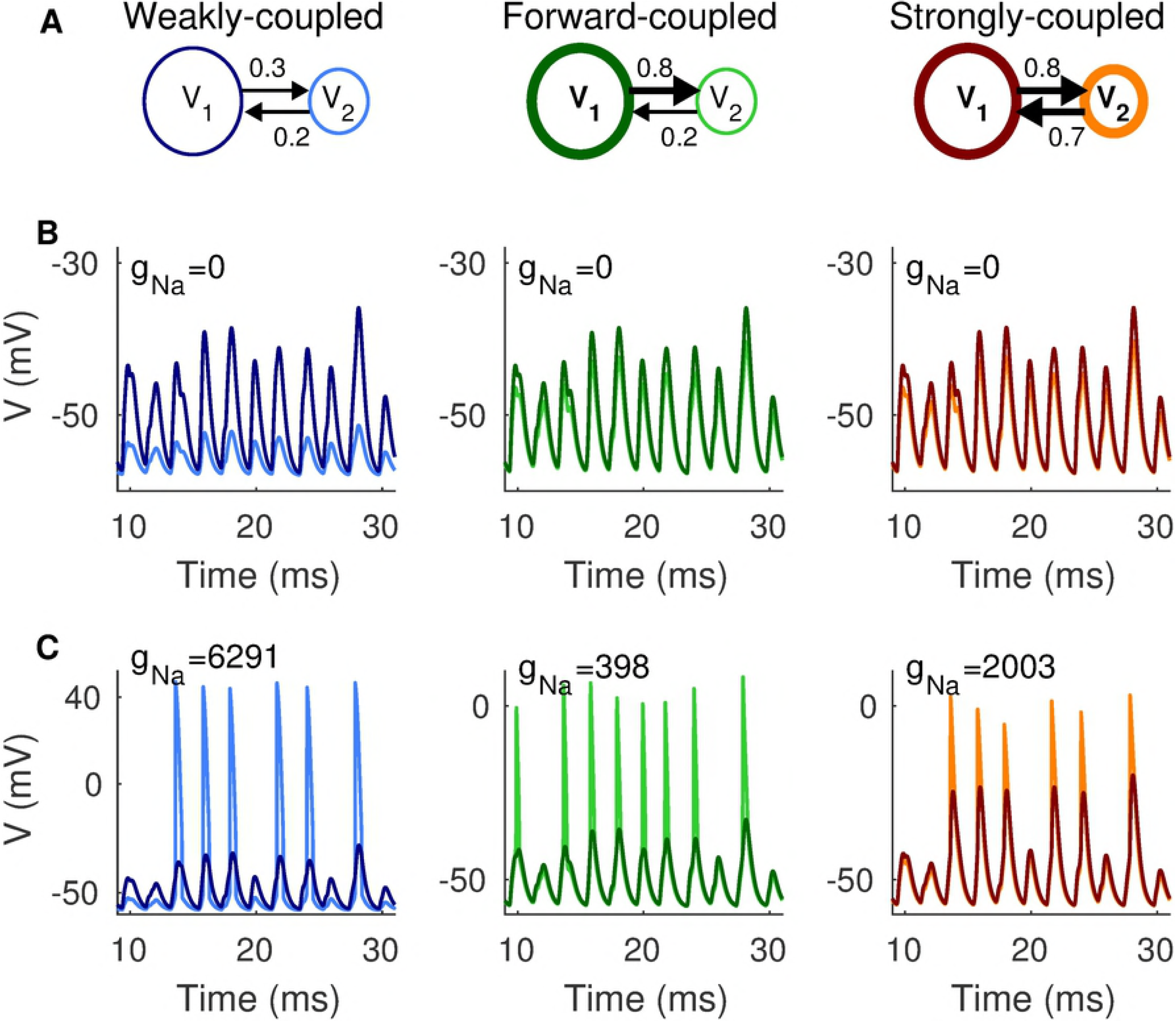
Responses of two-compartment models to 500 Hz coincident inputs. **A:** Schematic diagrams indicating coupling configurations for the weakly-coupled (left column), forward-coupled (middle column), and strongly-coupled (right column) models. **B:** Responses of passive models (*g*_*Na*_=0 nS). Darker colors indicate *V*_1_ (responses in Cpt1) and lighter colors indicate *V*_2_ (responses in Cpt2). *V*_1_ dynamics are nearly identical for all coupling configuration, as expected from our parameterization method. **C:** Responses of two-compartment models with 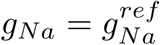. Occurrence and shape of spikes depends on coupling configuration, even though inputs are identical in all simulations. Notice the difference in y-axes; amplitude of *V*_2_ spikes is largest in the weakly-coupled model.

### Spiking dynamics depend on coupling configuration

We include spiking in the model by adding sodium current to Cpt2 (see Eq. 11). As described in Materials and methods, we define 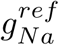 for each coupling configuration as the minimum level of *g*_*Na*_ at which two coincident inputs evoke a spike. We find it helpful to normalize *g*_*Na*_ to these reference values when comparing across coupling configurations. In Fig. 4C, we show responses to 500 Hz coincident inputs with *g*_*Na*_ set to 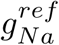. This results in *g*_*Na*_ = 6291 nS for the weakly-coupled model, *g*_*Na*_ = 398 nS for the forward-coupled model, and *g*_*Na*_ = 2003 nS for the strongly-coupled model. We do not include dynamic KLT current in these simulations. We use darker colors to show time-courses of voltage in Cpt1 (*V*_1_) and lighter colors to show time-courses of voltage in Cpt2 (*V*_2_).

These responses to identical inputs (same trains of synaptic inputs for each model, regardless of coupling configuration) reveal that spiking dynamics differ depending on coupling configuration. Coupling configuration dramatically changes the amplitude and shapes of spikes - the peaks of *V*_2_ in the weakly-coupled model are near 40 mV whereas the peaks of *V*_2_ in the forward-coupled and strongly-coupled models are near 0 mV (note the different axes). Spikes in the weakly-coupled model tend to be “all-or-none,” but the forward-coupled and strongly-coupled models can have more graded spike amplitudes. Not surprisingly, amplitudes of backpropagated spikes in Cpt1 depend on *κ*_2*→*1_. For models with weak backward coupling (the weakly-coupled and forward-coupled models), it is difficult to discern any changes in *V*_1_ due to spike generation in *V*_2_. For the strongly-coupled model, in contrast, *V*_1_ tracks *V*_2_ more closely and shows larger amplitude backpropagated spikes in Cpt1.

Importantly, the number and timing of spikes also changes with coupling configuration. In these traces, the forward-coupled model has two more spikes than the weakly-coupled and strongly-coupled models (see the “extra” spikes at 10 ms and 20 ms for the forward-coupled model). This anticipates our main result: coupling configuration affects spike generation and can alter the sensitivity of neurons to coincident inputs. If we view this example simulation using the analogy to signal detection theory and ROC analysis, described previously, we can say that the forward-coupled model correctly identifies two more coincident events (has two more “hits”) than the weakly-coupled and strongly-coupled models.

### Optimal coupling configuration for coincidence detection in two-compartment models with frozen KLT current

In a first set of simulations, we study the two-compartment model with passive subthreshold dynamics. The low-threshold potassium (KLT) current is “frozen” and included as part of the leak current (see Materials and methods). The only voltage-gated current in this set of simulations is the spike-generating sodium current in Cpt2. We quantify coincidence detection sensitivity by finding the maximum firing rate difference between coincident and non-coincident inputs for each coupling configuration (as described in Materials and methods). In Fig. 5, we report results for three stimulus frequencies (from left to right: 300 Hz, 500 Hz, and 700 Hz). We construct non-coincident inputs in two ways: in Fig. 5A we use out-of-phase sine wave inputs to the auditory nerve model. Time delays for out-of-phase inputs vary with frequency, so we also test coincidence detection sensitivity using sine wave inputs that are misaligned in time by a fixed 500 *µ*s time difference in Fig. 5B.

**Fig 5.**
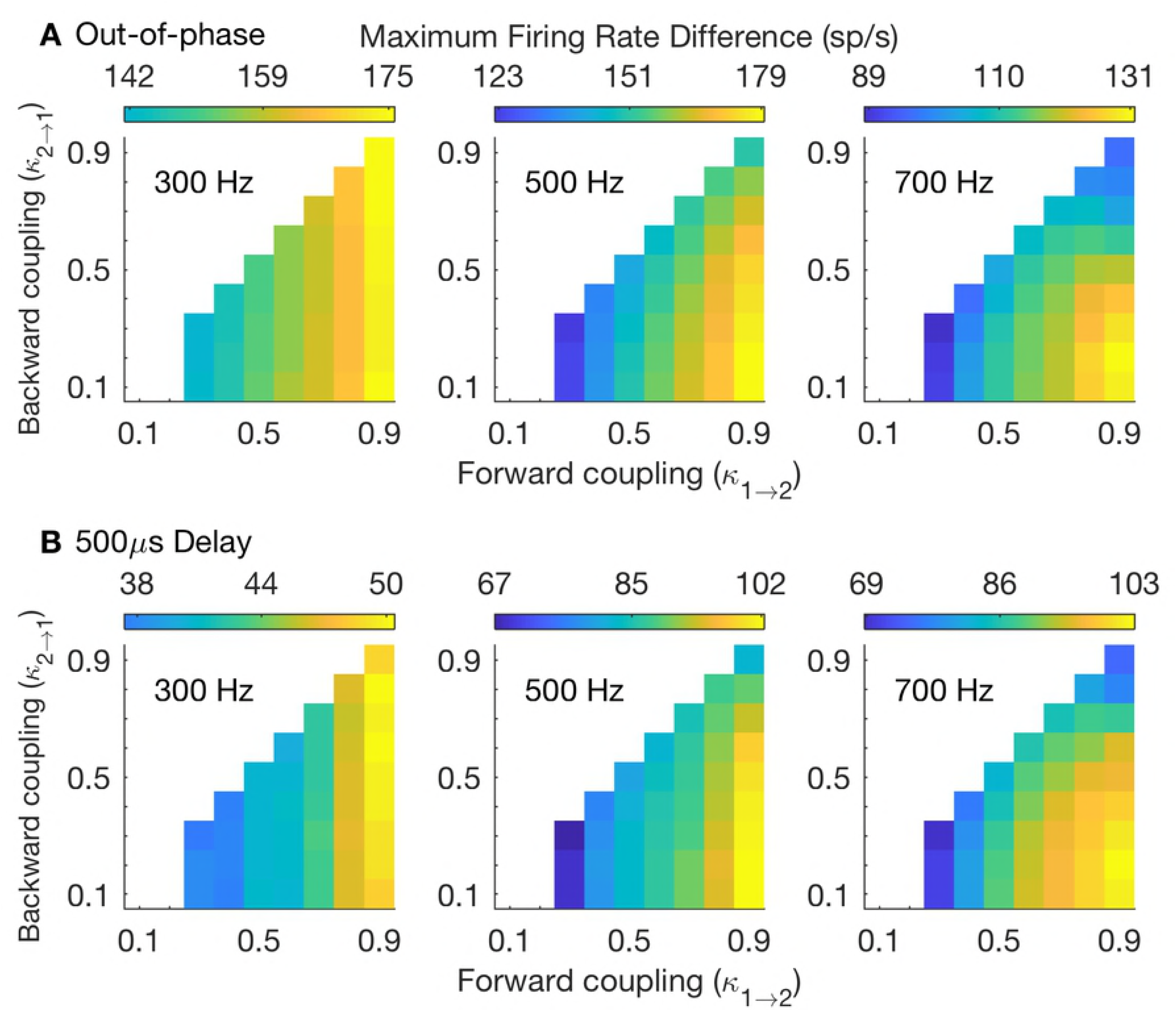
Coincidence detection sensitivity throughout parameter space of soma-axon coupling. **A:** Coincidence detection sensitivity measured as the maximum firing rate difference between responses to in-phase and out-of-phase stimuli. **B:** Coincidence detection sensitivity measured as the maximum firing rate difference between responses to in-phase stimuli and those with a 500 *µ*s time difference. In both rows: coincidence detection sensitivity is measured using three different stimulus frequencies (from left to right: 300 Hz, 500 Hz, 700 Hz). Color scheme in each panel is normalized so that the “lowest color” (dark blue) is the firing rate difference that is 65% of the highest value in that panel. In many panels these values are not reached and range of colors presented does not include these “low” blue colors.

Larger firing rate differences reflect better coincidence detection. In our analogy to a signal classification task, we say a large firing rate difference indicates a detector with a high “hit rate” in response to coincident inputs and a low “false alarm” rate in response to non-coincident events. We make two observations that we will explore in greater detail below. First, coincidence detection sensitivity improves with increases in forward coupling strength. That is to say, as one moves from left-to-right within each panel, the firing rate difference increases. Second, the combination of strong forward coupling (large *κ*_1*→*2_) and weak backward coupling (small *κ*_2*→*1_) enhances coincidence detection for high-frequency stimuli (notice the large firing rate differences in the lower right corner of panels for 500 Hz and 700 Hz stimuli).

In Fig. 6, we further detail how coincidence detection sensitivity changes with stimulus frequency. Strong forward coupling enhances coincidence detection for all frequencies above 200 Hz (the weakly-coupled model has the smallest firing rate difference across these frequencies). An advantage for the forward coupling configuration (combination of strong forward coupling and weak backward coupling) emerges for stimulus frequencies above about 400 Hz. These observations hold (qualitatively) regardless of whether we use out-of-phase stimuli for non-coincident inputs (Fig. 6A) or time-delayed stimuli (Fig. 6B). We provide tuning curves in S3 Fig for additional views of how coincidence detections sensitivity depends on stimulus frequency, *g*_*Na*_, and coupling configuration.

**Fig 6.**
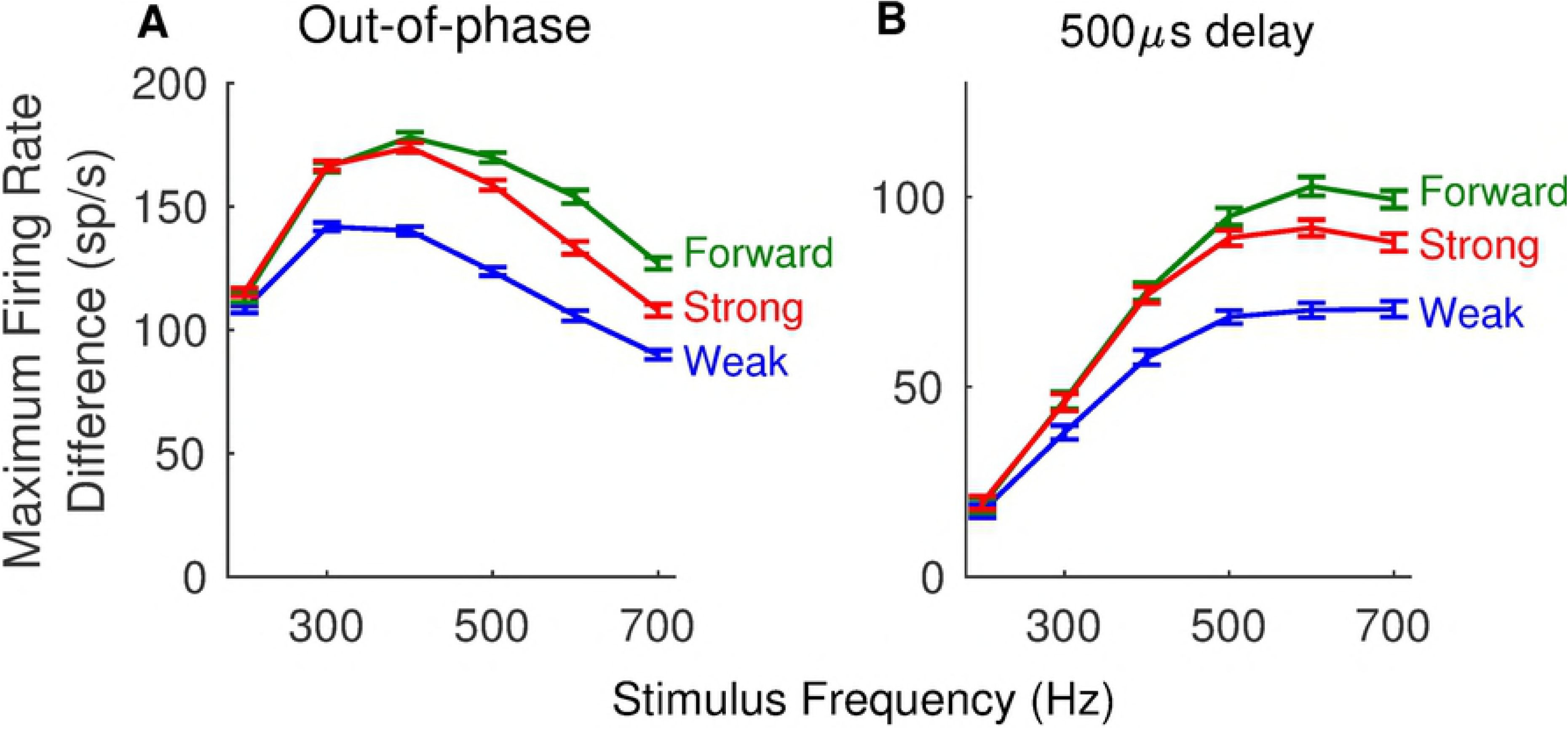
Dependence of coincidence detection sensitivity on stimulus frequency. A. Coincidence detection sensitivity measured as the maximum firing rate difference between responses to in-phase and out-of-phase stimuli. **B:** Coincidence detection sensitivity measured as the maximum firing rate difference between responses to in-phase stimuli and those with a 500 *µ*s time difference. In both panels: coincidence detection sensitivity is measured from responses to a range of frequencies (200 Hz to 700 Hz, in increments of 100 Hz), using the weakly-coupled (blue), forward-coupled (green), and strongly-coupled (red) models (see text, and Fig. 4 for definition of these models). Firing rates are computed from spikes counted in simulations with durations of 250 ms. Plotted values are the mean values computed from 100 repeated simulations, and error bars are standard errors in the means.

Why does strong forward coupling improve coincidence detection? And why does the specific combination of strong forward and weak backward coupling (the “forward-coupled” model) enhance high-frequency coincidence detection? We provide explanations below. First, we will demonstrate that strong forward coupling endows the two-compartment model with two properties that are advantageous for neural coincidence detection: phasic firing and sensitivity to input slope. Second, we will show that the specific combination of strong forward coupling and weak backward coupling shortens the refractory period of the two-compartment model. Neurons with short refractory periods can faithfully and rapidly respond to high-frequency sequences of coincident inputs. Neurons with longer refractory periods are disadvantaged when performing high-frequency coincidence detection, because they may “miss” opportunities to generate spikes in response to coincident inputs, and thus their “hit rate” (when thinking of these neurons as signal detectors) may be depressed.

### Strong forward coupling ensures phasic firing

Here we show that coupling configuration determines whether two-compartment models respond to steps of current injection with a single spike (phasic firing) or periodic spike trains (tonic firing). Phasic firing is advantageous for coincidence detection because it allows neurons to respond to rapid changes in stimuli – the concurrent arrival of several synaptic inputs, e.g. – and remain insensitive to slower changes in stimulus level that do not signal coincident inputs.

In these simulations we use direct current injection (not synaptic inputs) and frozen KLT current. See Fig. 7A for example waveforms of injected current. In Fig. 7B-D, we show time-courses of voltage in Cpt2 to these step current inputs, with *g*_*Na*_ set to 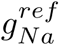. As the amplitude of the applied current increases, the weakly-coupled model exhibits a transition from quiescence (no firing) to periodic firing (Fig. 7B). The transition occurs via a Hopf bifurcation and is a tonic firing pattern. The forward-coupled and strongly-coupled models do not fire repetitively in these simulations. Instead, they transition from quiescence to a single spike elicited at the onset of the current injection (Fig. 7C and Fig. 7D). This response pattern is defined as phasic firing. We remind readers familiar with the Rothman-Manis [54] model that, in addition to introducing a two-compartment structure, we have also made changes to parameter values to reflect MSO physiology (time constant, input resistance, and resting potential). Thus, our observation of phasic firing in the strongly-coupled model does not contradict the observation by [36] of tonic firing in a Rothman-Manis-like model with frozen KLT, even though a one-compartment model can be viewed as an “extreme” case of a two-compartment model with *κ*_1*→*2_ = *κ*_2*→*1_ = 1.

**Fig 7.**
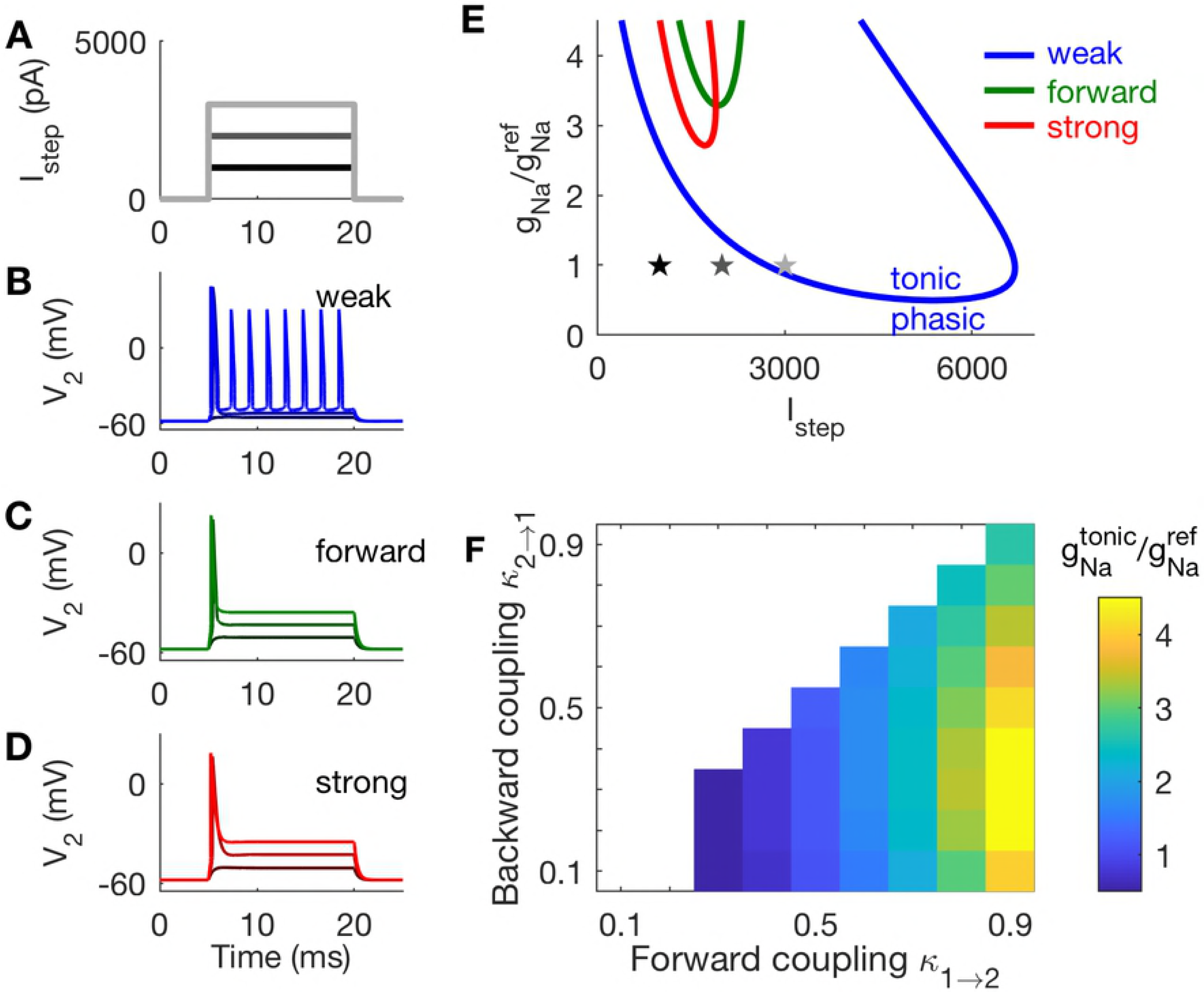
Responses of two-compartment models to constant current inputs. **A:** Waveforms of constant current steps, and *V*_2_ responses in panels **B-D**. **B:** Tonic firing in the weakly-coupled model. **C:** Phasic firing in the forward-coupled model. **D:** Phasic firing in the strongly-coupled model. **E:** Two-parameter bifurcation plot for the weak, forward, and strong coupling configurations. Models exhibit tonic firing for parameters within the U-shaped boundaries. The *g*_*Na*_ values on the y-axis are normalized to reference values 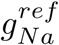. Stars indicate parameter values used in panels **A-D**. **F:** The lowest values of *g*_*Na*_ at which tonic firing is possible is 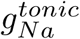. Models with strong forward coupling and weak backward coupling (lower right region in this panel) exhibit phasic firing over the largest range of (normalized) *g*_*Na*_ values.

We explore a range of *g*_*Na*_ values and trace out the boundary in parameter space between tonic and phasic firing. For each coupling configuration, we vary *g*_*Na*_ and the amplitude of the steady current (*I*_*step*_). We performed a two-parameter bifurcation analysis using the Auto feature in XPPAUT. The regions within the U-shaped curves in Fig. 7E are parameter combinations (*g*_*Na*_ and current levels) that produce repetitive firing.

The two-compartment model exhibits a tonic firing pattern if *g*_*Na*_ is set to a sufficiently large value. We identify the smallest values of *g*_*Na*_ at which repetitive firing to steady current can be observed (the lowest point on each U-shaped curve), and label these values 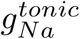. For example, the weakly-coupled model can exhibit tonic firing for values of *g*_*Na*_ larger than 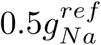, roughly. The forward-coupled and strongly-coupled models are phasic unless *g*_*Na*_ is larger than 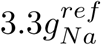 or 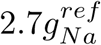, respectively. Values of the ratio 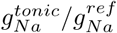 for all coupling configurations are shown in Fig. 7F. The value of this ratio increases with forward coupling strength. Phasic dynamics are closely associated with temporally-precise neural coincidence detection [34, 35]. This result indicates that phasic dynamics in models with stronger forward coupling are robust, in the sense that this coupling configuration can allow neurons to maintain phasic dynamics over a larger range of *g*_*Na*_ and input levels.

### Strong forward coupling enhances slope sensitivity

Another property associated with coincidence detection is slope sensitivity. Slope-sensitive neurons are those for which the rate of increase (slope) of an input – not just the input amplitude – can determine whether the neuron generates a spike [34]. As we now demonstrate, models with strong forward coupling are responsive to inputs with fast-rising inputs, whereas models with weak forward coupling can respond to inputs with slow-rising slopes. This is a second indication that strong forward coupling benefits neural coincidence detection.

We vary the slope of inputs using “ramps” of current, as shown in Fig. 8A. Time-courses of *V*_2_ In response to these three ramps are shown in Fig. 8B-D. We fix the maximum amplitude at 2500 pA for these inputs, and let the duration of the ramp vary with ramp slope.

**Fig 8.**
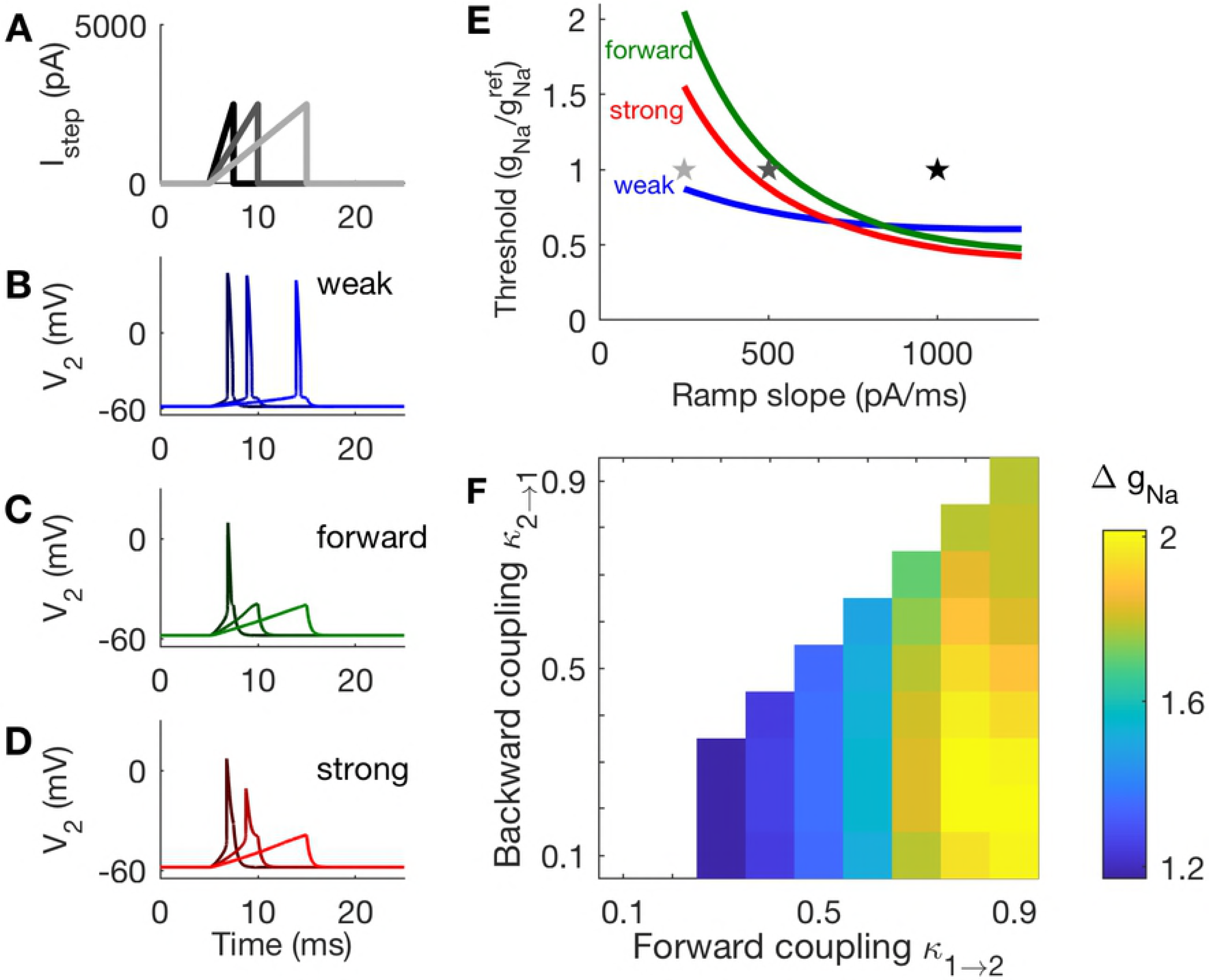
Responses of two-compartment models to ramped-current inputs. **A:** Waveforms of current ramps and *V*_2_ responses in panels **B-D**. **B:** The weakly-coupled model responds with a spike in response to all ramp inputs tested. **C:** The forward-coupled model responds with a spike in response to only the steepest ramp tested. **D:** The strongly-coupled model responds with a spike in response to the two ramps with steeper rising slopes. **E:** Sodium threshold is the smallest value of *g*_*Na*_ (plotted after normalization by 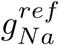) for which a ramp of a given slope elicits a spike. Threshold for the weakly-coupled model is relatively constant with ramp slope, but decreases with ramp slope for the forward-coupled and strongly-coupled models, indicating that these models are more slope-sensitive than the weakly-coupled model. Stars indicate parameter values used in panels **A-D**. **F:** Slope-sensitivity measured with Δ*g*_*Na*_, the ratio of threshold *g*_*Na*_ values computed from responses to input ramps with slopes 1000 pA/ms and 500 pA/ms. Larger values of Δ*g*_*Na*_ indicate coupling configurations that are more sensitive to the rising slope of inputs, and are found for models with strong forward coupling and weak backward coupling.

These simulations demonstrate that the forward-coupled model is the most slope-sensitive, as it only generates an action potential in response to the ramp with the steepest slope. In contrast, the weakly-coupled model spikes in response to all three ramps, and the strongly-coupled model spikes in response to two out of the three ramps.

We measure slope sensitivity by finding the smallest *g*_*Na*_ at which each model spikes in response to ramps of varying slopes (Fig. 8E). We again fix the amplitude of the current ramp at 2500 pA. The stars in Fig. 8E mark the parameter values used in Fig. 8B-D. For the weakly-coupled model, this measure of slope threshold is relatively constant as a function of ramp slope. In contrast, *g*_*Na*_ at firing threshold decreases dramatically for the forward-coupled and strongly-coupled models. This confirms that the forward-coupled and strongly-coupled models are more sensitive to input slopes than the weakly-coupled model. By setting *g*_*Na*_ appropriately, for example *g*_*Na*_ ≈ 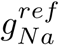, the forward-coupled and strongly-coupled models can be tuned to selectively spike in response to rapidly-rising inputs but not gradually-rising inputs.

To compare slope-sensitivity across all coupling configurations, we measured *g*_*Na*_ at firing thresholds in response to slow and fast rising ramps (500 pA/ms and 1000 pA/ms) and defined Δ*g*_*Na*_ to be the ratio of these values. A large value of Δ*g*_*Na*_ indicates that the model neuron can be made to fire selectively to steeply-rising ramps only (a slope-sensitive neuron), whereas values of Δ*g*_*Na*_ near one indicate that the model neuron responds similarly to ramps regardless of input slope. By this measure, models with strong forward coupling are more slope-sensitive than models with weak forward coupling. Moreover, the forward-coupled configuration (lower right corner of Fig. 8F) is optimal for slope-sensitivity.

### Comparison of models in the V_*2*_ – h phase plane

We gain an additional perspective on how spiking dynamics of the two-compartment model depend on coupling configuration by examining the *V*_2_ – *h* phase plane. For the weak, forward, and strong coupling configurations, we plot *h* and *V*_2_-nullclines in Fig. 9. The *h*-nullcline, defined by the function *h* = *h*_*∞*_(*V*_2_), is shown in black in each panel and is the same for all coupling configurations. Nullclines for *V*_2_ are shown with colored lines in each panel. We calculate these nullclines by setting *V*_1_ to fixed values: -58 mV (*V*_*rest*_), -40 mV, and -30 mV. These curve are “sections” (at the selected *V*_1_ values) through the “null surfaces” of the full three-dimensional *V*_1_ – *V*_2_ – *h* phase space. The rationale for computing *V*_2_-nullclines for certain fixed values of *V*_1_ is that *V*_1_ can be viewed as the “input” to Cpt2 (recall Eq. 1). Notice that the *V*_2_-nullclines are truncated in these figures – a “right branch” at higher values of *V*_2_ in which the *V*_2_-nullcline is an increasing function of *V*_2_ is not pictured, and the local maximum (“left knee”) of the *V*_2_-nullclines with *V*_1_ = –58 mV are at larger values of *h* than what is pictured in Fig. 9.

**Fig 9.**
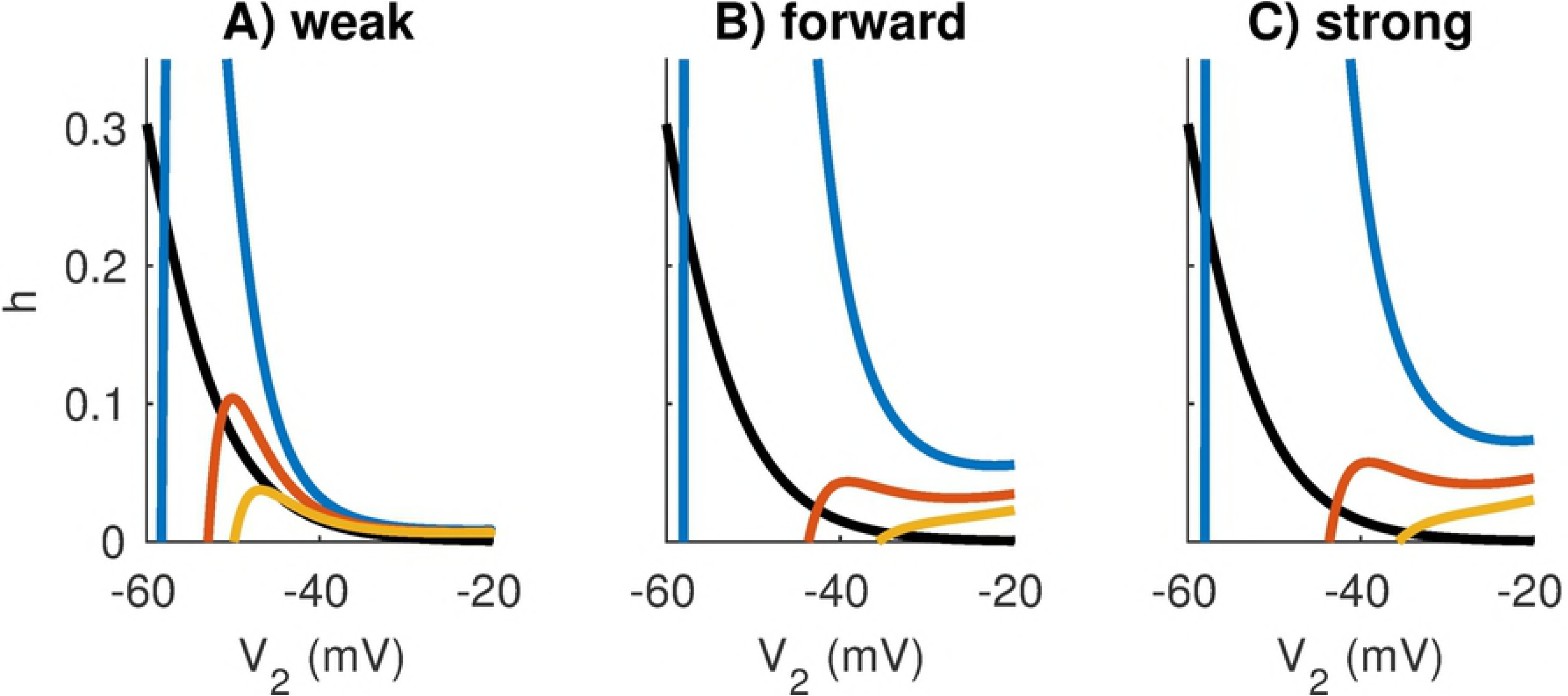
Nullclines in the *V*_2_-*h* plane for fixed values of *V*_1_. The *V*_2_-nullclines (colored curves) are sections of three-dimensional nullsurfaces at fixed values of *V*_1_: the resting value (*V*_1_ = -58 mV for blue curve) and two depolarized values (*V*_1_ = -40 mV for red curve, *V*_1_ = -30 mV for orange curve). The *h*-nullcline is identical in all models (black curve) and given by *h* = *h*_*∞*_(*V*_2_). Coupling configurations are, from left to right, **A:** weakly-coupled, **B:** forward-coupled, and **C:** strongly-coupled. Notice that *V*_1_-dependent fixed point moves from the left to middle branch of the *V*_2_-nullcline for the weakly-coupled model. This marks the transition to tonic firing that is absent in the forward-coupled and strongly-coupled models. The *V*_2_-nullclines have “cubic” shapes as is typical for excitable dynamics, but we only exhibit a small portion of the *V*_2_ and *h* axes.

We use these phase plane diagrams to illustrate the distinction between tonic firing in the weakly-coupled model and phasic firing in the strongly-coupled and forward-coupled models. For all models, the *V*_2_-nullclines shift rightward as *V*_1_ increases. In the case of weak coupling, the intersection between the *h* and *V*_2_-nullclines (the *V*_1_-dependent fixed point of the system) crosses over from the left branch of the *V*_2_-nullcline to the middle branch as these nullclines shift rightward. This signals the destabilization of the fixed point via a Hopf bifurcation as *V*_1_ increases beyond a threshold value. For the forward-coupled and strongly-coupled models, by contrast, the left knee (local maximum) of the *V*_2_-nullcline disappears as *V*_1_ increases. For these models, then, dynamics of the spike-generator cease to be excitable for large *V*_1_ values. This prevents the possibility of repetitive (tonic) firing to steady inputs.

These phase plane diagrams also reveal the role sodium inactivation plays in enhancing slope-sensitivity in models with strong forward coupling. In Fig. 9, we observe that increases in *V*_1_ cause larger rightward shifts in the *V*_2_-nullcline for the strongly-coupled and forward-coupled model, as compared to the weakly-coupled model. As a result, for identical *V*_1_ values, the fixed points in the *V*_2_ – *h* plane are located at larger *V*_2_ values and smaller *h* values for the models with stronger forward coupling. By construction, there is less attenuation from Cpt1 to Cpt2 in the models with stronger forward coupling. Thus, a given depolarization of *V*_1_ produces larger depolarizations of *V*_2_ in these models compared to the weakly-coupled model. Since larger depolarizations of *V*_2_ cause greater inactivation of sodium current (reducing the *h* gating variable), inputs that elicit spikes in the strongly-coupled and forward-coupled models must depolarize the cell rapidly to activate sodium current on a timescale faster than inactivation of sodium by the *h* gating variable. In sum, slope-sensitivity in this model arises from the dynamic, voltage-gated, negative feedback of sodium inactivation, and models with stronger forward coupling are more effective at engaging this process.

### Combination of strong forward coupling and weak backward coupling shortens the refractory period

We have discussed phasic firing and slope-sensitivity as factors that make models with strong forward coupling more precise coincidence detectors than models with weak forward coupling. We have not yet, however, established why the particular combination of strong forward coupling and weak backward coupling is advantageous for high-frequency coincidence detection. To do this, we investigate refractory periods and post-spike recovery dynamics. Specifically, we show that models that have both strong forward coupling and weak backward coupling have short refractory periods and rapid recovery after a spike.

The responses in Fig. 10 illustrate how refractory period changes with coupling configuration. The stimuli for these simulations are a sequence of two EPSG events (imagine, for example, a brief input from each of the two ears). The amplitude of each EPSG event is three times the size of a unitary synaptic input from the auditory nerve model. The first EPSG in the sequence evokes a spike and, depending on the time delay and the coupling configuration, the trailing EPSG may or may not evoke a spike (even though it is the same amplitude as the first EPSG). In the simulations shown, the weakly-coupled model does not produce a second spike for a time delay of 2 ms, but the forward-coupled and strongly-coupled models do (Fig. 10B-D). In all these simulations, *g*_*Na*_ was set 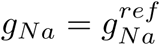.

**Fig 10.**
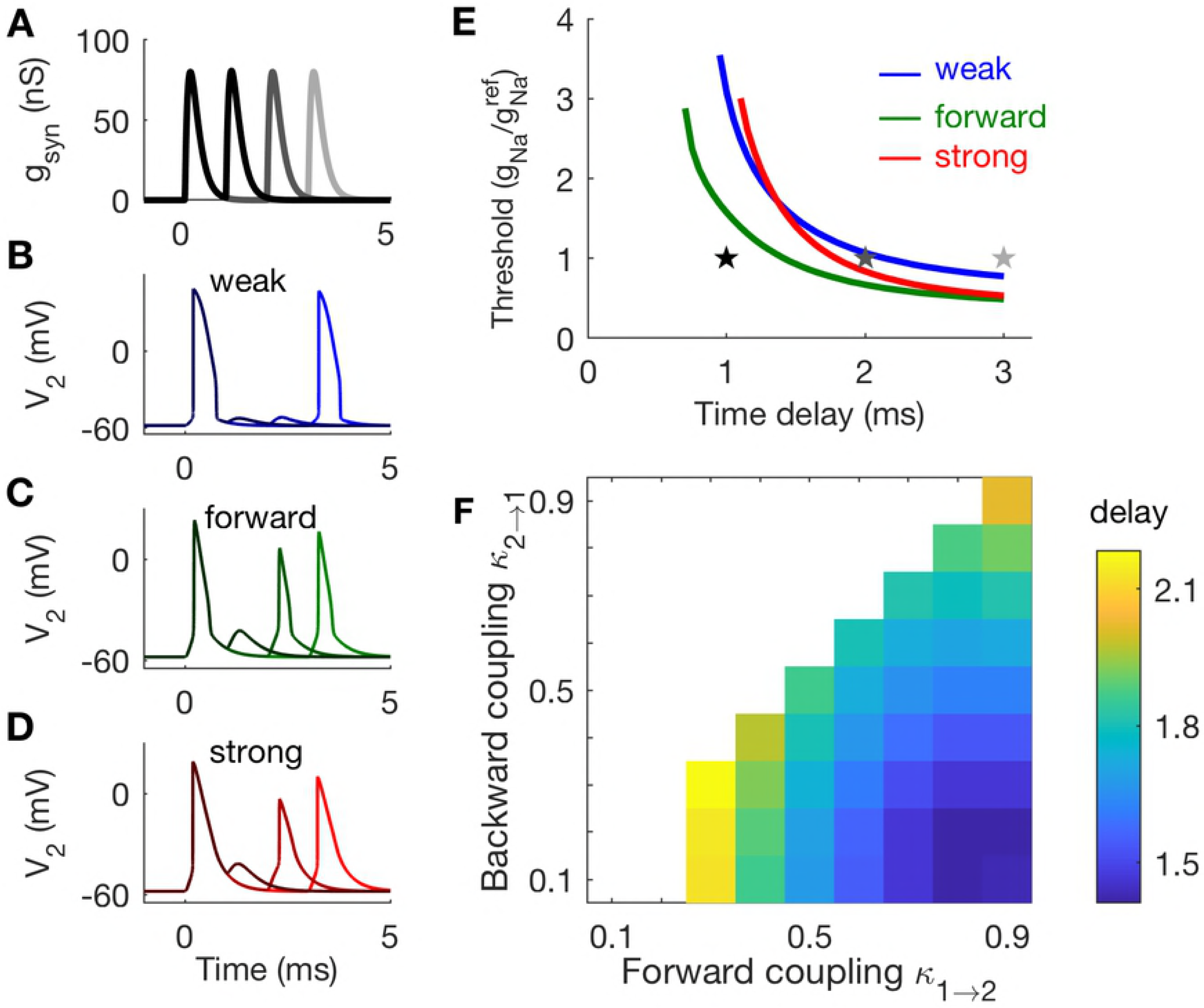
Responses of two-compartment models to pairs of synaptic excitatory inputs. **A:** Waveforms of excitatory post-synaptic conductances (EPSGs) and *V*_2_ responses in panels **B-D**. Inputs consists of an EPSGs at *t* = 0 and a second EPSG at a later time. **B:** The weakly-coupled model spikes in response to trailing EPSG delayed by 3 ms. **C:** The forward-coupled model spikes in response to trailing EPSGs delayed by 2 ms or 3 ms. **D:** The strongly-coupled model spikes in response to trailing EPSGs delayed by 2 ms or 3 ms. **E:** Sodium threshold is the smallest value of *g*_*Na*_ (plotted after normalization by 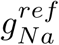) for which both EPSGs elicit spikes for a given slope time delay. The smallest time delay at which two spikes can be elicited is the refractory period (left-most point of each curve). Stars indicate parameter values used in panels **A-D**. **F:** Refractory period is measured by finding the minimum time delay at which the model neuron fires in response to both EPSG events (using *g*_*Na*_ = 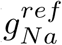). The coupling configurations with shortest refractory periods (smallest delay thresholds) are those with strong forward coupling and weak backward coupling.

To further investigate the effect of coupling configuration on refractory period for these three models, we determined the smallest *g*_*Na*_ value for which both EPSGs produced spikes, for varying time delays (Fig. 10E). The amplitude of each EPSG in the pair of inputs was three times the amplitude of a unitary auditory nerve fiber input (same as in Fig. 10A). This calculation identifies the refractory period – the smallest time delay for which the neuron produces a spike in response to both inputs. Refractory periods in these simulations are 0.7 ms for the forward-coupled model, 0.95 ms for the weakly-coupled model, and 1.1 ms for the strongly-coupled model. In addition to having the smallest refractory period among these three models, the forward-coupled model also has the lowest thresholds across all time-delays. This indicates that the forward-coupled does not require excessive sodium conductance to respond rapidly to successive inputs.

To probe the refractory period across all coupling configurations, we fixed *g*_*Na*_ at reference values, and measured the smallest time delay for which the model neuron could respond to both EPSGs. By this measure, models with strong forward coupling and weak backward coupling have the smallest refractory period (lower right corner of Fig. 10F).

We explain the effect of coupling configuration on refractory period by examining post-spike recovery for the weakly-coupled, forward-coupled, and strongly-coupled models in the *V*_2_ – *h* phase plane (Fig. 11). The monotonically decreasing (black) curve is the *h*-nullcline and the cubic curves (only partially shown) are the *V*_2_-nullclines for *V*_1_ fixed at rest and at -42 mV. These are similar to the nullclines shown in Fig. 9. The thin (green) curves show trajectories of the response of each model in the *V*_2_ –*h* phase plane to a pair of synaptic events separated by a 1.5 ms time delay. Sodium conductance is set to 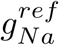 and the first synaptic event elicits a spike. Responses of the models to the trailing inputs depend on refractory period. For these parameter sets, only the forward-coupled model can respond to both inputs in the pair of successive synaptic inputs.

**Fig 11.**
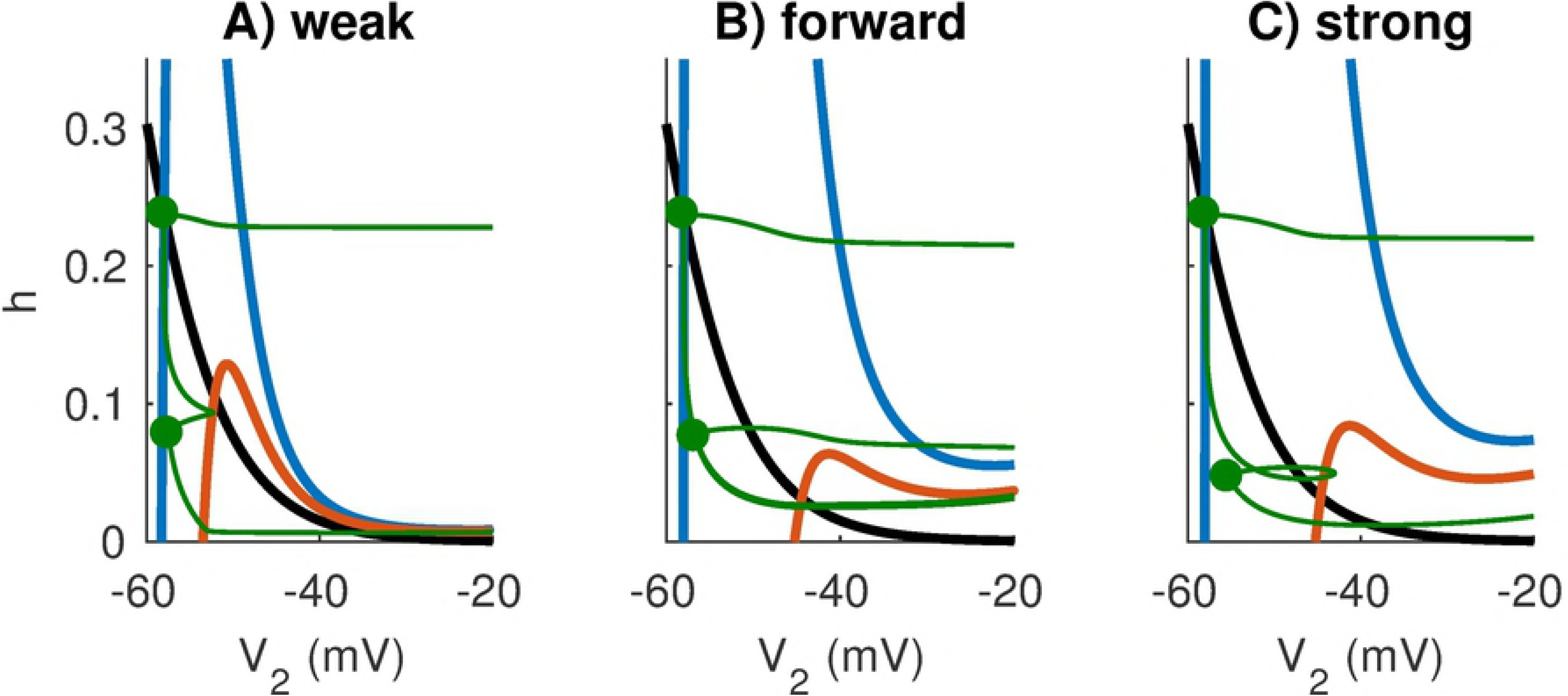
Nullclines in the *V*_2_**-***h* plane for fixed values of *V*_1_ and trajectory of response to pair of EPSGs. The *V*_2_-nullclines (colored curves) are sections of three-dimensional nullsurfaces at fixed values of *V*_1_: the resting value (*V*_1_ = -58 mV for blue curve) and a depolarized value of *V*_1_ = –42 mV (red curve) chosen to roughly represent the effect of an EPSG input with amplitude three times that of a unitary event from the auditory nerve model. The *h*-nullcline is identical in all models (black curve) and given by *h* = *h*_*∞*_(*V*_2_). Coupling configurations are, from left to right, **A:** weakly-coupled, **B:** forward-coupled, and **C:** strongly-coupled. In these simulations we set *g*_*Na*_ = 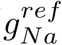. The green trajectory is the response of the model to a pair of EPSGs with time delay of 1.5 ms and amplitude three times that of a unitary auditory nerve fiber input. The green dots identify the onsets of each EPSG event. Only the forward-coupled model fires in response to both events in these simulations (**B**). The forward-coupled model has the shortest refractory period because excitatory inputs recruit sodium activation to decreases the height of the left knee of the *V*_2_-nullcline, and the post-spike dynamics of this model allow sodium inactivation (*h* gating variable) to recover rapidly.

To gain a qualitative sense of why the forward-coupled model spikes in response to both inputs (and therefore has a shorter refractory period), we compare the state of the neuron as it evolves through the *V*_2_ – *h* phase plane to the height of the left knee of the *V*_2_-nullcline. As *V*_1_ increases in response to an excitatory input, the *V*_2_-nullcline shifts downward (transition from blue to red curves). More specifically, the height of the left knee of the *V*_2_-nullcline shifts downward as *V*_1_ increases. If an excitatory input (which depolarizes *V*_1_) shifts the left knee of the *V*_2_-nullcline below the position of the trajectory in *V*_2_ – *h* phase plane, then the *V*_2_ variable will move (quickly, on a fast time-scale) to the right branch of the *V*_2_-nullcline. This represents the spike upstroke, and is not visible in its entirety in Fig. 11 because the right branch of the *V*_2_-nullcline is outside the “field of view” of these figures. In these phase plane figures, we see that the second excitatory input, which depolarizes *V*_1_ to roughly -42 mV, only elicits a spike in the forward-coupled model.

There are two factors that contribute to the short refractory period in the the forward coupling model. First, observe that the height of the left knee of the *V*_2_-nullclines are lower for the models with strong forward coupling (forward-coupled and the strongly-coupled models) than for the weakly-coupled model. In models with strong forward coupling, an increase in *V*_1_ propagates to *V*_2_ with minimal attenuation. This increase in *V*_2_ activates sodium current which lowers the height of the left knee of the *V*_2_-nullcline. As a consequence, models with strong forward coupling are more excitable (more responsive to an excitatory input) during the recovery period than models with weak forward coupling.

Second, observe that *h* (the sodium inactivation gating variable) recovers more quickly in models with weak backward coupling. To see this, compare the height of the lower green dot across the models (lower dots represent the state of the neuron 1.5 ms after the onset of the first synaptic input). The gating variable has the value *h* = 0.08 for the weakly-coupled and forward-coupled models at that instant, whereas *h* = 0.05 for the strongly-coupled model. Sodium inactivation recovers more slowly in the strongly-coupled model. Spike recovery is slow in the strongly-coupled model because the spike in Cpt2 depolarizes *V*_1_ (due to the strong backward coupling), which then prevents *V*_2_ from rapidly returning to rest (due to the strong forward coupling). In models with weak backward coupling, by contrast, *V*_1_ does not depolarize substantially during a spike, and thus Cpt1 can act as a current sink to help return *V*_2_ to rest.

These differences in sodium inactivation recovery are subtle, but the combination of the two factors discussed above gives the forward-coupled model a double advantage in responding to high-frequency inputs. Excitation in Cpt1 transfers to Cpt2 efficiently to depolarize the spike-generator, and the spike-generator “resets” quickly (via recovery of the *h* gating variable) to accommodate rapid generation of action potentials.

### Coincidence detection sensitivity in two-compartment models with dynamic KLT current

In the preceding sections, we have detailed the advantages of strong forward coupling generally, and weak backward coupling for high-frequency stimuli, for coincidence detection sensitivity in a two-compartment neuron model. With the exception of the spike-generating sodium current, the two-compartment model we have considered to this point has been passive. We questioned how our findings would change if additional voltage-gated currents were included. Of particular interest in the context of neural coincidence detection in the MSO is the low threshold potassium (KLT) current. This current is prominent in MSO neurons and enhances their coincidence detection sensitivity [33]. We therefore repeated our test of coincidence detection sensitivity with dynamic KLT conductance (see Materials and methods).

In Fig. 12 we show coincidence detection sensitivity measured from responses to three input frequencies (from left to right: 300 Hz, 500 Hz, and 700 Hz). In the top row, 10% of the total conductance in Cpt1 at rest is dynamic KLT conductance. In the bottom row, 10% of the total conductance in Cpt2 at rest is dynamic KLT conductance. The format of each panel is similar to Fig. 5 with the color scale in each panel representing the maximal firing rate difference between in-phase and out-of-phase inputs.

**Fig 12.**
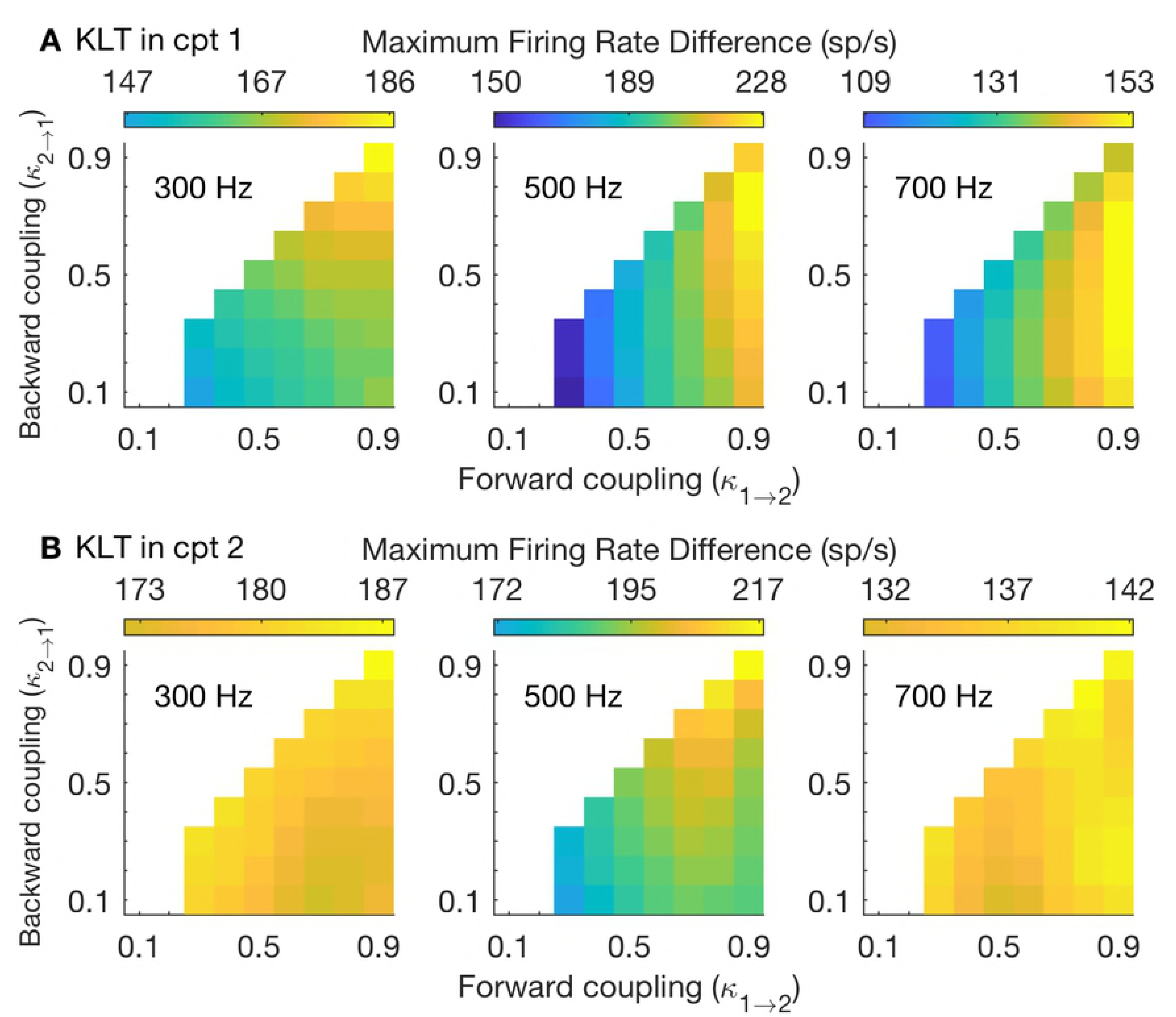
Coincidence detection sensitivity in two-compartment models with dynamic KLT. Coincidence detection sensitivity is measured using three different stimulus frequencies (from left to right: 300 Hz, 500 Hz, 700 Hz). Color scheme in each panel is normalized so that the “lowest color” (dark blue) is the firing rate difference that is 65% of the highest value in that panel. In many panels these values are not reached and range of colors presented does not include these “low” blue colors. **A:** 10% of leak conductance in Cpt1 replaced with voltage-gated, low-threshold potassium (KLT) conductance. **B:** 10% of leak conductance in Cpt2 replaced with voltage-gated, low-threshold potassium (KLT) conductance.

Upon comparing to our previous results using a passive model (frozen KLT), we observe some differences. Dynamic KLT in the input region (Cpt1, top row) improves coincidence detection sensitivity for all model configurations. While models with strong forward coupling and weak backward coupling (lower right corner of each panel) remain as effective coincidence detectors, the optimal configuration shifts to models with stronger backward coupling. For the 300 Hz stimulus, for instance, the largest firing rate differences are achieved for models with strong forward and strong backward coupling (upper right corner). Models with strong backward coupling can more effectively “make use” of the KLT current because *V*_2_ spikes propagate back into Cpt1 to activate the KLT current.

Dynamic KLT in the output region (Cpt2, bottom row) also improves coincidence detection sensitivity for all model configurations, but the greatest increases are in models with weak coupling. As a result, coincidence detection sensitivity is nearly uniform across all model configurations, especially in responses to lower and higher frequency inputs. Dynamic KLT in Cpt2 tends to provide the most benefit for models with weak coupling because it provides a secondary source of voltage-gated, dynamic, negative feedback in these models for which sodium inactivation does not suffice to establish dynamics conducive to coincidence detection, including phasic responses to steady inputs and slope-sensitivity (recall Fig. 7 and Fig. 8).

We compare coincidence detection sensitivity across a range of stimulus frequencies for models with the weak, forward, and strong coupling configurations and that include dynamic KLT, see Fig. 13. As above, we test models with dynamic KLT conductance in Cpt1 (Fig. 13A1 and B1), and models with dynamic KLT conductance in Cpt2 (Fig. 13A2 and B2). Results for models with dynamic KLT are shown in thick lines. For reference, we also include our earlier results using the frozen KLT model (thin lines, same as results shown in Fig. 6). The results are consistent with our observations from Fig. 12. In particular, we find that dynamic KLT conductance improves coincidence detection sensitivity (relative to the passive model) for all coupling configurations and nearly all stimulus frequencies. Moreover, the benefit of KLT depends on coupling configuration. Dynamic KLT added to Cpt1 (soma) improves coincidence detection sensitivity the most for the strongly-coupled model. Dynamic KLT added to Cpt2 (axon) improves coincidence detection sensitivity the most for the weakly-coupled model.

**Fig 13.**
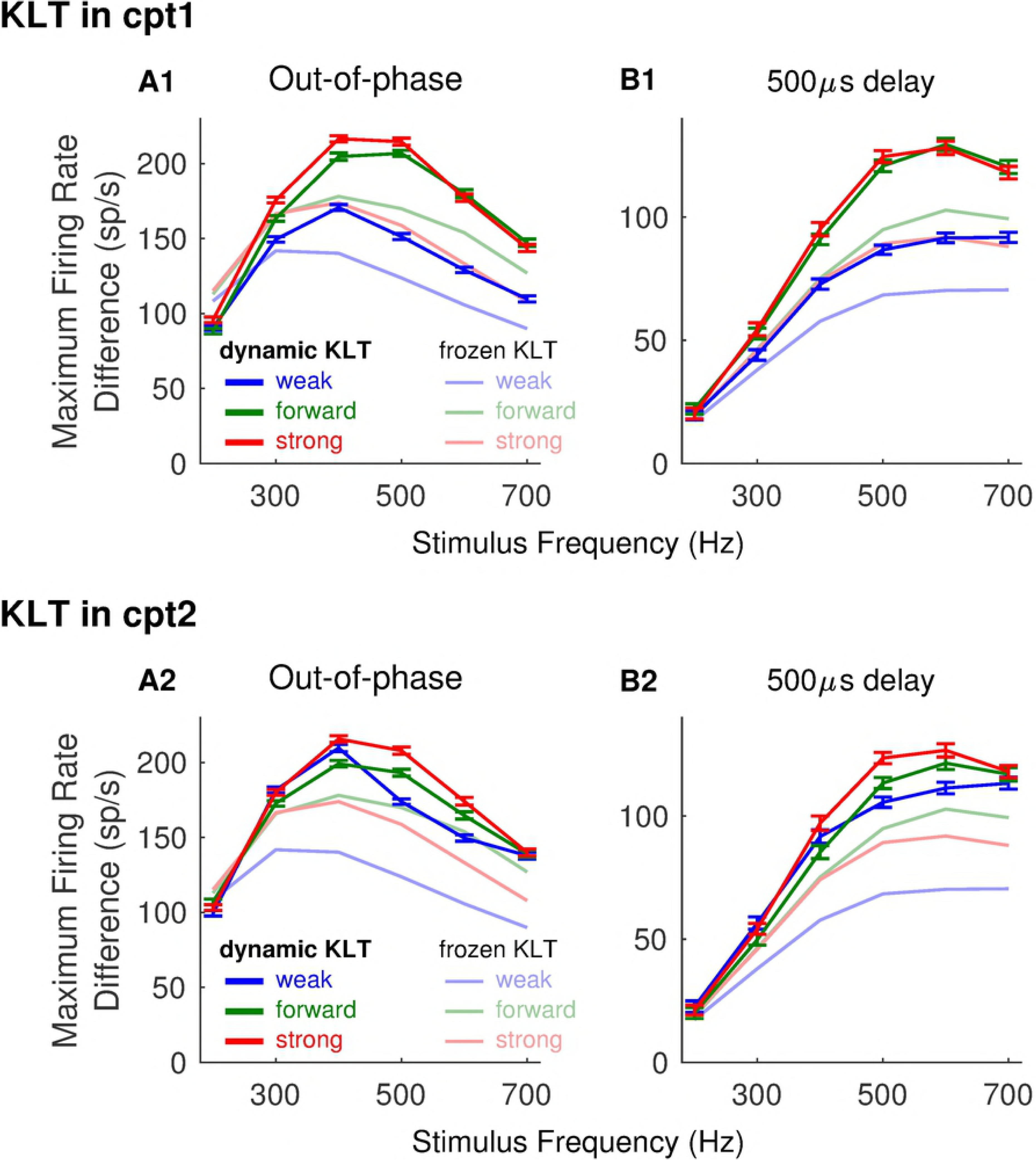
Dependence of coincidence detection sensitivity on stimulus frequency for models including dynamic KLT. Firing rate differences for models with dynamic KLT are shown in thick lines. For reference, we also show firing rate differences for models with frozen KLT as thin lines (frozen KLT model, same as Fig. 6A). Top row (**A1** and **B1**): 10% of resting conductance in Cpt1 (input region) is dynamic KLT conductance. Bottom row (**A2** and **B2**): 10% of resting conductance in Cpt2 (output region) is dynamic KLT conductance. Left column (**A1** and **A2**): Coincidence detection sensitivity measured as the maximum firing rate difference between responses to in-phase and out-of-phase stimuli. Right column (**B1** and **B2**): Coincidence detection sensitivity measured as the maximum firing rate difference between responses to in-phase stimuli and stimuli with 500 *µ*s time difference. In all panels: coincidence detection sensitivity is measured from responses to a range of frequencies (200 Hz to 700 Hz, in increments of 100 Hz), using the weakly-coupled (blue), forward-coupled (green), and strongly-coupled (red) models. Firing rates are computed from spikes counted in simulations with durations of 250 ms. Plotted values are the mean values computed from 100 repeated simulations, and error bars are standard errors in the means.

## Discussion

We systematically examined how soma-to-axon coupling affects neural coincidence detection. We characterized coupling configuration by two parameters (*κ*_1*→*2_ and *κ*_2*→*1_) representing strength of coupling between “input” region (soma and dendrite) and “output” region of a cell (axon and axon initial segment). We also identified a separation of time scales between the “slow” subthreshold dynamics in the input region and the “fast” dynamics in the spike generator. We measured coincidence detection sensitivity in the model neurons by viewing them as “observers” of their own synaptic inputs performign a signal detection task — we interpreted spiking in response to coincident inputs as a measure of correct detection (“hits”) and spiking responses to non-coincident inputs as a measure of incorrect detection (“false positives”). Combining these analyses enabled us to elucidate how coupling configuration (described by a few parameters) affects coincidence detection properties of the two-compartment model neuron.

We fixed parameter values based on known properties of principal cells of the medial superior olive (MSO). These neurons are among the first “binaural” neurons in the mammalian auditory brainstem. MSO neurons receive inputs that originate from both ears and respond preferentially to coincident inputs. The remarkable temporal sensitivity of these coincidence detector neurons is critical for sound localization processing in mammals.

### Strong soma-to-axon coupling enhances neural coincidence detection

Specializations that support temporally-precise coincidence detection in MSO neurons include voltage-gated currents active at membrane potentials near resting values [53], fast and well-timed excitatory synapses [27], and dendritic structure (bipolar dendrites, that segregate inputs from opposite ears onto opposite dendrites) [29, 31], see also [25] for review. In this work, we showed that soma-axon coupling is an additional structural specialization that can enhance neural coincidence detection.

By performing a thorough search through the space of coupling configuration, we found that strong forward (soma-to-axon) coupling improved coincidence detection sensitivity. And, moreover, the asymmetric “forward-coupled” configuration of strong forward coupling and weak backward coupling was the optimal configuration for coincidence detection in response to higher frequency inputs (500 Hz to 700 Hz). We identified advantages of strong forward coupling for neural coincidence detection including phasic and slope-sensitive spiking dynamics, and (for the forward-coupled model) short refractory periods. These advantages depended on the action of the sodium inactivation gating variable, which is the sole source of voltage-gated, dynamic, negative feedback in the version of the two-compartment model with frozen KLT.

### Dynamic KLT current further enhances neural coincidence detection

KLT current is an additional source of negative feedback and one known to be prominent in MSO neurons in soma and dendrites regions [31], as well as axon regions [38]. We found that dynamic KLT current improved coincidence detection for nearly all coupling configuration and stimulus frequencies. There were notable interactions between coupling configuration and KLT current. Dynamic KLT current improved coincidence detection in neurons with strong forward and backward coupling so that this “strongly-coupled” configuration (the configuration most similar to a one-compartment model) became optimal for coincidence detection in response to intermediate-frequency stimuli. In addition, dynamic KLT current localized to Cpt2 (the spike-generator region of the neuron model), could “rescue” coincidence detection sensitivity in neurons with weak soma-axon coupling so that this “weakly-coupled” configuration could exhibit coincidence detection sensitivity on par with other models. There was some loss of efficiency when dynamic KLT current was included (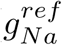 values increased by 5% to 30%), but these differences are modest compared to the order of magnitude differences in 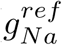 across coupling configurations.

The picture that emerges is one in which structural and dynamical features can combine to provide neurons with a suite of specializations that are useful for temporally-precise coincidence detection. Strong forward coupling appears to be a “natural” configuration for coincidence detection. Moreover, the specific combination of strong forward coupling and weak backward coupling is advantageous for high frequency coincidence detection. The benefits of these coupling configurations can be supplemented with appropriately-targeted KLT current. The need for multiple, complementary mechanisms that enhance coincidence detection MSO neurons has been explored previously, but usually for one-compartment models (Na inactivation and KLT activation as two sources of negative feedback [36]; KLT and h currents as currents that regulate input resistance [32]). The effect of KLT is considerable and well-studied, so here we emphasized the role of structure (soma-to-axon coupling). The “intrinsic” advantage of the forward-coupling configuration may help maintain coincidence detection sensitivity in scenarios in which KLT is less effective (for instance, KLT current can be inactivated on a slow time-scale not considered in our model). Other features of MSO neurons are specialized for coincidence detection including inhibitory synaptic inputs [57, 62, 63] and dendritic structure [29–31]. The variety of physiological “tools” MSO neurons use to perform coincidence detection emphasizes the exceptional nature of the temporally-precise computations these neurons perform.

We tested coincidence detection sensitivity with out-of-phase inputs and inputs with a fixed 500 *µ*s delay. We observed qualitatively similar results for both types of inputs. The latter stimulus may be more relevant in studying neural coincidence in the context of sound localization. Time differences in this context would be created by differences in travel times of sounds arriving at the two ears (“interaural time differences”) and are limited by animal head size. In humans, for instance, maximal interaural time differences created by head size are approximately 700 *µ*s.

### MSO neurons appear “forward-coupled”

Our model is phenomenological – the two-compartments are “lumped” representations of input and output regions. We do not, therefore, resolve structural details of dendrites or spike initiation zones (see [38] for an example of the latter). Nonetheless, we can make qualitative observations that relate our findings to MSO physiology.

Action potentials in the MSO are likely generated in a spike initiation zone near the soma, and back-propagated action potentials in the soma are small and graded [37]. This indicates a “strict electrical segregation of the soma and dendrites from the axonal initiation zone,” (in the words of [37]). In the context of our model, this corresponds to weak backward coupling (small value of *κ*_2*→*1_). Backpropagation of signals into the dendrites is further attenuated due to the low input resistance of these neurons and the strong effects of voltage-gated potassium current [31]. Additionally, current injection into the soma reliably evokes action potentials that propagate into the axon [37]. This suggests a configuration in which the soma has a strong effect on the spike-generator (minimal attenuation, large value of *κ*_1*→*2_ in our model).

Taken together then, it appears that MSO neurons may be structured in a “forward-coupled” manner, consistent with our observations that this configuration confers advantages for coincidence detection by engaging sodium inactivation as dynamic negative-feedback mechanism, by promoting rapid “resetting” of the spike generator (shortening the refractory period, which enables high-frequency spiking), and by enabling efficient spike generation (smaller sodium conductance required).

The complete picture of MSO excitability and axonal structure is doubtless more complicated. Recent computational simulations provide evidence that spike generation may occur throughout MSO axons (initial segment and multipole nodes of Ranvier) [38]. Spike generation at more distal locations on the axon can preserve excitability in response to high-frequency stimuli by preventing inactivation of sodium channels [38, 44]. Studies of coincidence detector neurons in related structures in the avian auditory show that excitability of these neurons can be adjusted via modulation of ion channel density in spike generator regions [42–44]. This raises intriguing questions about plasticity in the spike initiation zone, and dynamic regulation of the soma-to-axon connection.

### A framework for investigating neural structure, dynamics, and computation

We have formulated a family of two-compartment models to investigate neural coincidence detection in MSO neurons. We showed that parameters in this two-compartment framework can be chosen in a principled manner to explore the range of coupling configurations, while maintaining similar passive dynamics in the input region. With this approach, we identified how structure (the nature of soma-axon coupling) affected dynamics in the spike-generator region, and, in turn, how these differences in dynamics affect the sensitivity of coincidence detector neurons to synaptic inputs.

Our approach provides a unifying view of structure and function in neurons performing an identified computation. It is one that should find applications in studies of other neurons. Coincidence detector neurons in the auditory brainstem of owls, for instance, have been modeled as two-compartment structures [41]. The two-compartment idealization has also been useful for investigating dynamics of bursting [45, 49, 72], bistability [46], oscillations [47], and resonance [50] in neurons, and could also describe signaling between a (large) dendrite region and a (small) dendritic spine. Our framework for creating, and systematically exploring, a parameter space of soma-axon coupling configurations, can be used to shed further light on the relationship between structure, dynamics, and function in these and other neural systems.

## Supporting information

**S1 Fig. Reference** *g*_*Na*_ **for models with 10% dynamic low-threshold potassium (KLT) conductance. A:** Reference *g*_*Na*_ across parameter space of coupling strengths, for model without dynamic KLT. This panel reproduces Fig. 2B, see text for definition of reference *g*_*Na*_. **B:** Reference *g*_*Na*_ for two-compartment models with 10% of leak conductance in Cpt1 replaced by dynamic KLT conductance. **C:** Reference *g*_*Na*_ for two-compartment models with 10% of leak conductance in Cpt2 replaced by dynamic KLT conductance.

**S2 Fig. Effect of stimulus frequency and coupling configuration on best Na conductance.** The value of *g*_*Na*_ at which the two-compartment model achieves its best coincidence detection sensitivity (maximal firing rate difference between response to in-phase and out-of-phase inputs) for (**A**) 300 *Hz* stimuli, (**B**) 500 *Hz* stimuli, and (**C**) 700 *Hz* stimuli. **D:** Detailed view of these “best” *g*_*Na*_ values for the weakly-coupled, forward-coupled, and strongly-coupled models as function of input frequency.

**S3 Fig. Coincidence detection tuning curves for the weak, forward, and strong coupling models.** Firing rates in response to 300 Hz (left column), 500 Hz (middle column) and 700 Hz (right column) stimuli. Sodium conductance values are selected so that the firing rate for coincident inputs (0 ms time difference) for each model is 150 spikes/second (top row), 250 spikes/second (middle row), or 350 spikes/second (bottom row).

**S1 File. Excel spreadsheet of figure data.**

## Acknowledgments

Large-scale computations were performed using the Ohio Supercomputer Center (OSC). JHG thanks the OSC for a start-up allocation of computing time.

